# Phosphoproteome modifications and cortical circuit dysfunction are linked to the early-stage progression of alpha-synuclein aggregation

**DOI:** 10.1101/2025.01.24.634820

**Authors:** Sayan Dutta, Jennifer A. Hensel, Alicia N. Scott, Rodrigo Mohallem, Leigh-Ana M. Rossitto, Hammad F. Khan, Luke A. Diehl, Teshawn Johnson, Bryce D. Colón, Jacob E. Gold, Christina R. Ferreira, Jackeline F. Marmolejo, Xu Chen, Krishna Jayant, Uma K. Aryal, Laura Volpicelli-Daley, Jean-Christophe Rochet

**Affiliations:** Borch Department of Medicinal Chemistry and Molecular Pharmacology, Purdue University, West Lafayette, IN, 47907, USA; Department of Comparative Pathobiology, College of Veterinary Medicine, Purdue University, West Lafayette, Indiana, 47907, USA; Department of Neurosciences, School of Medicine, University of California, San Diego, 92161, USA; Weldon School of Biomedical Engineering, West Lafayette, Indiana, 47907, USA; Metabolite Profiling Facility, Bindley Bioscience Center, Purdue University, West Lafayette, IN 47907; Center for Neurodegeneration and Experimental Therapeutics, University of Alabama at Birmingham, Birmingham, AL 35294, USA; Purdue Institute for Integrative Neuroscience, Purdue University, West Lafayette, IN, 47907, USA; Purdue Proteomics Facility, Bindley Bioscience Center, Purdue University, West Lafayette, Indiana, 47907, USA; Division of Biology and Biological Engineering, California Institute of technology, CA, 91125, USA; Department of Biochemistry and Molecular Biology, Oklahoma State University, Stillwater, OK, 74078, USA

## Abstract

Cortical dysfunction is increasingly recognized as a major contributor to the non-motor symptoms associated with Parkinson’s disease (PD) and other synucleinopathies. Although functional alterations in cortical circuits have been observed in preclinical PD models, the underlying molecular mechanisms are unclear. To bridge this knowledge gap, we investigated tissue-level changes in the cortex of rats and mice treated with alpha-synuclein (aSyn) seeds using a multi-omics approach. Our study revealed significant phosphoproteomic changes, but not global proteomic or lipid profiling changes, in the rat sensorimotor cortex 3 months after intra-striatal injection with aSyn preformed fibrils (PFFs). Gene ontology analysis of the phosphoproteomic data revealed that PFF administration impacted pathways related to synaptic transmission and cytoskeletal organization. Similar phosphoproteomic perturbations were observed in the sensorimotor cortex of mice injected intrastriatally or intracortically with aSyn PFFs. Functional analyses demonstrated increased neuronal firing rates and enhanced spike-spike coherence in the sensorimotor cortex of PFF-treated mice, suggesting that aSyn seeds induced cortical circuit dysfunction. Bioinformatics analysis of the altered phosphosites indicated the involvement of several kinases, including casein kinase-2 (CK2) and Protein Kinase A (PKA). Enrichment of kinase-activating pT/pY MAPK (ERK) phosphosites revealed apparent engagement of the ERK signaling cascade and led to the identification of pERK1/2-positive intraneuronal granulovacuolar body (GVB)-like structures specifically in aggregate-bearing neurons. Collectively, these findings highlight the importance of phosphorylation-mediated signaling pathways in the cortical response to aSyn pathology spread in PD and related synucleinopathies, setting the stage for developing new therapeutic strategies.

## Introduction

Parkinson’s disease (PD) is a neurodegenerative disorder involving a complex array of motor and nonmotor symptoms (Russell, Drozdova et al. 2014, Jellinger 2018). A key pathological feature of PD is the death of dopaminergic neurons that project from the substantia nigra pars compacta (SNpc) to the striatum, a degenerative process underlying the disease’s motor symptoms (Shulman, De Jager et al. 2011). PD neuropathology is also characterized by the presence in multiple brain areas of intracellular Lewy pathology enriched with fibrillar forms of the presynaptic protein, aSyn (Spillantini, Crowther et al. 1998, Crowther, Daniel et al. 2000). Lewy pathology, consisting of both Lewy bodies and Lewy neurites, is found in the brains of individuals with PD as well as other neurodegenerative diseases associated with pathological aSyn aggregation, such as dementia with Lewy bodies (DLB) (Spillantini, Crowther et al. 1998) and multiple system atrophy (MSA) (Schweighauser, Shi et al. 2020), collectively known as synucleinopathies. Although current therapies effectively manage PD motor symptoms for several years, their efficacy declines over time, and no effective treatments exist to address non-motor symptoms or slow disease progression. (Pirooznia, Rosenthal et al. 2021).

Post-mortem analyses of PD brains at various stages of the disease indicate that aSyn aggregates initially appear in the olfactory bulb and brainstem and progressively spread to the midbrain and cortex as the disease advances (Braak, Del Tredici et al. 2003, Beach, Adler et al. 2009). Additionally, the injection of preformed fibrils (PFFs) of recombinant aSyn in rodent striatum leads to the formation of neuronal Lewy-like pathology through anatomically connected pathways (Luk, Kehm et al. 2012, Paumier, Luk et al. 2015, Henderson, Cornblath et al. 2019). aSyn pathology propagation in both synucleinopathy patients and PFF-injected rodents is believed to involve the release of aggregates from affected neurons into the extracellular space. The secreted aggregates are then taken up by neighboring healthy neurons, where they promote the aggregation of endogenous cytosolic aSyn via a seeding mechanism(Peng, Trojanowski et al. 2020). aSyn aggregates secreted into the CSF can be detected as a readout of underlying aSyn pathology in patients’ brains using seed amplification assays (Siderowf, Concha-Marambio et al. 2023).

aSyn is involved in the regulation of synaptic vesicle release and membrane dynamics as part of its normal physiological function (Runwal and Edwards 2021, Sharma and Burre 2023). Conversely, aSyn aggregates are known to have various pathogenic effects, including mitochondrial impairment and the dysregulation of proteostasis and lipid homeostasis (Wang, Becker et al. 2019, Fanning, Selkoe et al. 2021, Flores-Leon and Outeiro 2023, Zalon, Quiriconi et al. 2024). Primary hippocampal neurons treated with aSyn PFFs show evidence of altered synaptic activity and impaired functional network connectivity (Volpicelli-Daley, Luk et al. 2011, Froula, Henderson et al. 2018). In addition, electrophysiological analyses of brain slices from PFF-treated mice have revealed that seeded aSyn aggregation impairs neurotransmission (Chen, Nagaraja et al. 2022). Accordingly, intracellular seeded aSyn propagation is predicted to lead to various forms of cellular dysfunction in rodent PFF models, involving either a loss of aSyn normal function as the monomeric protein is recruited into growing aggregates (Chen, Nagaraja et al. 2022, Calabresi, Di Lazzaro et al. 2023), or a gain of toxic function due to the accumulation of the aggregates themselves (Calabresi, Mechelli et al. 2023). However, information is lacking regarding the functional perturbations associated with aSyn pathology propagation in the brains of PFF-injected rodents, especially during the early stages of aggregate spread that occur well before the onset of neurodegeneration.

In this study, we carried out proteomic, phosphoproteomic, and lipid profiling of cortical homogenates from rats three months after intrastriatal injection with either aSyn PFFs or control aSyn monomer. This timeframe allowed for the formation of Lewy-like aggregates in PFF-treated animals without resulting in overt neurodegeneration or behavioral deficits. Phosphoproteomic analysis revealed pronounced PFF-dependent changes, leading to the identification of multiple phosphoprotein hits involved in cellular transport and synaptic function. These phosphoproteomic perturbations were validated in a mouse aSyn PFF model, which exhibited corresponding PFF-dependent changes in neurocircuitry function. Many of the phosphoproteins up-regulated in the brains of PFF-treated rats or mice were predicted substrates of various kinases, including casein kinase-2 (CK2) and protein kinase A (PKA). The phosphoproteomic data also revealed evidence of PFF-dependent enrichment of activating pT/pY ERK1/2 phosphosites, suggesting potential ERK pathway engagement. Immunohistochemical (IHC) validation of PFF-dependent ERK activation led to the discovery of unique pERK1/2-positive granulovacuolar bodies (GVB) in aggregate-bearing cortical neurons, signifying early-stage stress responses and altered proteostasis. Collectively, these results provide insights into cellular perturbations that could accompany the spread of cortical aSyn pathology in the brains of individuals with PD and other synucleinopathy disorders.

## Results

### PFF-induced aSyn aggregation leads to phosphoproteomic (but not global proteomic) changes in rat sensorimotor cortex

To identify molecular perturbations associated with the propagation of aSyn pathology, rats were injected bilaterally in the striatum with aSyn preformed fibrils (PFFs) or monomer (**Figure 1A,B**) using established methods (Paumier, Luk et al. 2015). Brain samples were collected 3 months post-injection and processed for histology or proteomic, phosphoproteomic, or lipid profiling. IHC analysis revealed that aSyn PFFs, but not the control monomer, induced the formation of aggregates that stained positive for pS129-aSyn in various brain regions, including the sensorimotor cortex (**Figure 1C; Supplementary Figure 1A,B**). Aggregates resembling Lewy bodies or Lewy neurites were found to be distributed among all cortical layers, especially layers IV and V, based on an analysis of sections stained for N-terminal EF-hand calcium-binding protein 1 (NECAB1), a neuronal protein specifically expressed in cortical layers IV/V (**Figure 1C,D**).

**Figure 1:**
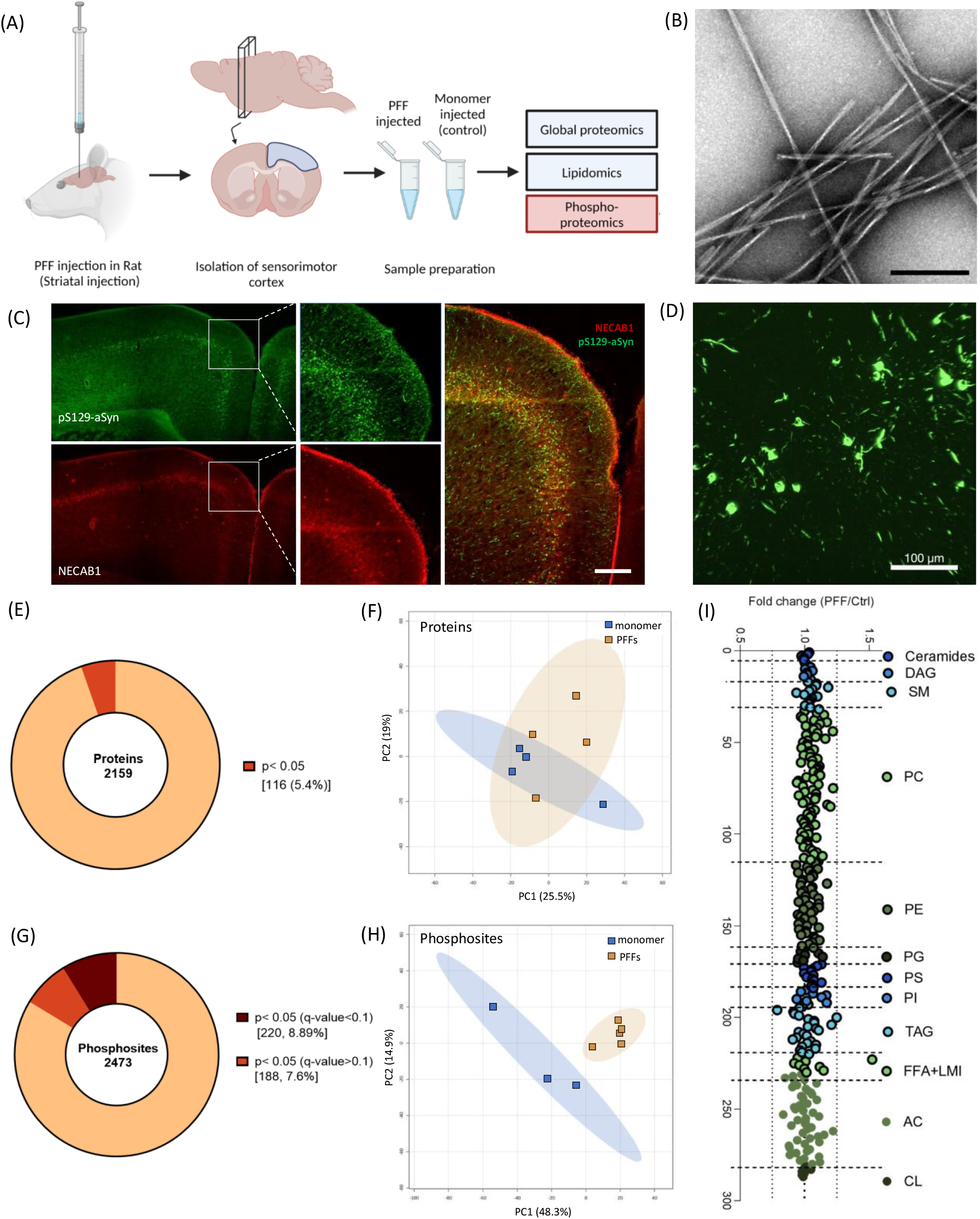
Rats injected intrastriatally with aSyn PFFs show evidence of phosphoproteomic changes in the sensorimotor cortex. (A) Overview of the multi-omics approach used to identify cortical changes associated with aSyn aggregation in the rat PFF model (created using https://BioRender.com) (B) Representative images of aSyn preformed fibrils (PFFs) visualized by transmission electron microscopy. Scale bar: 100 nm. (C) Representative images of a rat cortical section stained for pS129-aSyn (81a, green) and the cortical layer IV/Layer V-specific marker NECAB1 (red) 3 months after aSyn PFF injection in the striatum (n = 3 animals). Scale bar: 200 µm. (D) Higher magnification image of the cortex from a PFF-injected rat showing pS129-aSyn aggregates resembling Lewy bodies and Lewy neurites. (E) Pie chart representation of protein hits obtained via global proteomic analysis of homogenates prepared from rat sensorimotor cortex 3 months after intrastriatal injection with aSyn PFFs or monomer. The chart shows the percentage of hits with p≥0.05 or p<0.05. Each hit was detected in ≥70% of samples in at least one experimental group. (F) Graph showing the results of unassigned/unsupervised PCA of the log_2_-transformed intensities of all protein hits identified in the cortical homogenates described in (E). (G) Pie chart representation of unique phosphosite hits obtained via phosphoproteomic analysis of the cortical homogenates described in (E). The chart shows the percentage of hits with p<0.05 (q<0.1) or p<0.05 (q>0.1). Each hit was detected in ≥70% of samples in at least one experimental group. (H) Graph showing the results of unassigned/unsupervised PCA of the log_2_-transformed intensities of all phosphosite hits identified in the cortical homogenates described in (E). (I) Graph showing lipid profile perturbations (expressed as fold change in lipid level) in the cortical homogenates described in (E). None of the changes were below the significance threshold of p<0.05. DAG, diacylglycerols; SM, sphingomyelin; PC, phosphatidylcholine; PE, phosphatidylethanolamine; PG, phosphatidylglycerol; PS, phosphatidylserine; PI, phosphatidylinositol; TAG, triacylglycerols; FFA+LMI, free fatty acids and lipid metabolism intermediates; AC, acylcarnitines; CL, cardiolipin.

Analysis of cortical homogenates via liquid chromatography-tandem mass spectrometry (LC-MS/MS) led to the identification of 31,772 unique peptides (label-free quantitation (LFQ) > 0 in at least one sample) derived from 2,799 unique proteins. Following rigorous selection criteria (intensity observed in >70% of the samples in at least one of the two treatment groups), 2,159 proteins met this stringent cutoff, with only 116 proteins exhibiting a p-value below the cutoff value for significance (p<0.05; **Figure 1E**). Unsupervised 2D principal component analysis (PCA) revealed two independent overlapping clusters (**Figure 1F**), confirming that there were no pronounced global proteomic changes attributable to cortical aSyn aggregation at the 3-month time point. Next, we conducted similar analyses of other PD-relevant brain regions including the amygdala and SN (**Supplementary Figure 1C-I**). Consistent with our findings from studies of the sensorimotor cortex, no significant proteomic differences were observed in these regions when comparing the PFF- and monomer-injected animals.

Based on evidence that aSyn pathology can alter kinase localization or activity (Ryu, Kim et al. 2008, Garcia-Yague, Lastres-Becker et al. 2021), we next examined phosphoproteomic changes in sensorimotor cortex samples from rats injected with aSyn PFFs or monomer. Changes in the phosphorylation states of synaptic proteins have been implicated as key cellular mechanisms in the dynamic regulation of neuronal synapses, even in the absence of altered protein expression levels (Kohansal-Nodehi, Chua et al. 2016, Li, Wilkinson et al. 2016, Engholm-Keller, Waardenberg et al. 2019). We detected 3,565 unique phosphopeptides (LFQ>1) from 1,509 distinct proteins. After data filtering, 2,473 phosphosites with distinct multiplicities, mapping to 1058 proteins, were detected. Of these phosphosites, 220 (i.e., 8.9%, corresponding to 161 unique proteins) exhibited significant changes in abundance (q-value<0.1) in the brains of PFF-treated rats compared to animals injected with aSyn monomer (**Figure 1G**). Unsupervised PCA revealed two distinct clusters, providing evidence of differences in the phosphoproteome profile between these groups (**Figure 1H**). These findings suggest that seeded aSyn aggregation induces cell signaling perturbations resulting in altered phosphorylation events.

Finally, given the well-documented interaction of aSyn in both monomeric and aggregate form with cellular membranes (Fanning, Selkoe et al. 2021, Runwal and Edwards 2021, Flores-Leon and Outeiro 2023, Sharma and Burre 2023), as well as evidence of changes in lipid homeostasis in the brains of PD patients (Battis, Xiang et al. 2023), we profiled 1,687 lipid species belonging to 13 lipid classes in cortical homogenates from aSyn PFF- and monomer-treated rats (**Supplementary Figure 2A**, **Supplementary Table 1**). Of these, 287 lipids exceeded the detection threshold (≥1.3-fold over blank measurements; **Supplementary Figure 2B**, **Supplementary Table 1**). However, a comparison between the two groups revealed no substantial changes in lipid composition. With the exception of a single lipid species, all changes were within ± 25% of control values (**Figure 1I**, **Supplementary Figure 2C**). Consequently, none of the lipids met our predefined criteria for significance (|FC| ≥ 2), and, therefore, they were not included in downstream ontology and pathway analyses. Similar lipidomic analyses were performed on homogenates prepared from amygdala and SN, yielding comparable results with no significant differences between groups (data not shown).

### Gene ontology analysis reveals alterations in cellular processes involved in synaptic transmission in the brains of PFF-injected rats

Among the 220 significantly up-or down-regulated phosphosites (i.e., with q<0.1), a 2-fold increase or decrease in intensity was observed in 172 and 9 phosphosites, respectively, in cortical homogenates prepared from rats injected with aSyn PFFs versus monomer (**Figure 2A,B**). A cluster heatmap of log_2_-transformed intensities for the 50 lowest q-value phosphosites further highlights distinct patterns of up- and-down-regulated phosphoproteins in samples obtained from PFF-versus monomer-treated animals (**Figure 2C**). Only a subset of phosphosites within individual phosphoproteins exhibited PFF-dependent dysregulation, suggesting that seeded aSyn aggregation alters site-specific phosphorylation rather than overall protein expression in the rat PFF model. This conclusion is also supported by the absence of any significant changes in abundance of the parent peptides in the global proteomics dataset.

**Figure 2:**
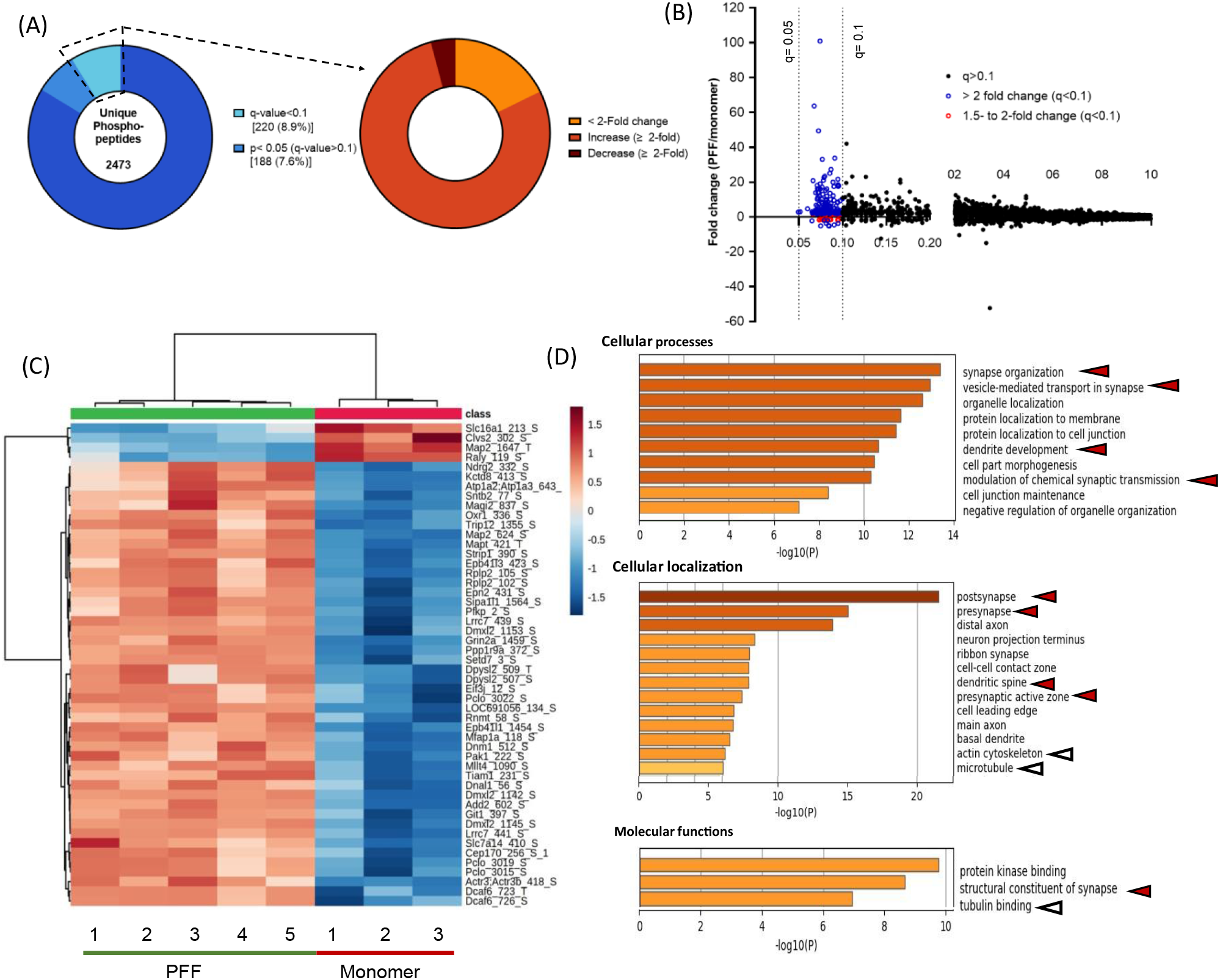
Gene ontology analysis of the top up- or down-regulated phosphoproteins in the sensorimotor cortex of PFF-treated rats reveals perturbations of cellular structures and functions involved in synaptic transmission. (A) Pie chart representations of phosphosite hits obtained via phosphoproteomic analysis of homogenates prepared from rat sensorimotor cortex 3 months after intrastriatal injection with aSyn PFFs or monomer. The chart on the left is identical to that shown in Figure 1H. The chart on the right shows the percentage of phosphosite hits with q<0.1 that exhibit a fold change of <2 (yellow) or are up- or down-regulated by a factor of ≥2 (orange and brown, respectively). Each hit was detected in ≥70% of samples in at least one experimental group. (B) Plot showing the phosphosite distribution represented as intensity fold change (PFF/monomer) versus q-value. Phosphosites with q<0.1 that are up- or down-regulated by a factor of 1.5 to 2 or >2 are represented by red and blue symbols, respectively. (C) Cluster heatmap showing the z-scored log_2_-transformed intensities of the top 50 up- or down-regulated phosphosites (i.e., phosphosites with the lowest q-values) in the cortical homogenates described in (A). Red and blue colors correspond to an increase or decrease (respectively) in phosphosite levels in rats injected with PFFs versus monomer, and the color intensity represents the Z-score-normalized log_2_(intensity) value. Peptide names are listed as ‘protein name_phosphoresidue number’. (D) Charts showing enriched GO terms within cellular processes (top), cellular localization (middle), and molecular functions (bottom) that represent groups of phosphoproteins in the sensorimotor cortex impacted by PFF treatment. GO analysis was carried out on proteins containing phosphosites with q<0.1 and a fold change of ≥2 (see panel A). Arrowheads highlight items relevant to synaptic transmission (red) or the cytoskeleton (white).

Genes associated with significantly up- and down-regulated phosphosites were further examined via gene ontology (GO) enrichment analysis. ‘Synaptic organization’ emerged as the most enriched cellular processes term (**Figure 2D, top**), involving >25 genes encoding pre- and post-synaptic scaffolding proteins (e.g., Piccolo, Bassoon, Rims1, Shank1, Shank3), proteins involved in modulating synaptic vesicle pools and neurotransmitter release (e.g., Dmxl2, Sv2A), and proteins linked to the generation of action potentials (e.g., Grin2a, Grin2b). Other highly ranked cellular processes terms included ‘vesicle-mediated transport in synapse’, ‘dendrite development’, and ‘modulation of chemical synaptic transmission,’ suggesting that the observed phosphoproteomic changes reflected alterations in synaptic function associated with PFF-mediated aSyn aggregation. Examination of enriched cellular localization terms revealed ‘post-synapse’ (39 genes) and ‘pre-synapse’ (32 genes) as the two most enriched terms (Figure 2D, middle), with minimal overlap. Pathways related to cytoskeletal organization were also identified from our analysis of enriched ‘cellular localization’ and ‘molecular functions’ terms (Figure 2D, middle and bottom). Moreover, many phosphosites exhibiting changes in abundance in the brains of PFF-treated rats were derived from kinases or proteins associated with kinases, suggesting that PFF administration induced changes in kinase activity. Notably, the largest cluster within the ‘molecular functions’ category included 25 ontology terms related to ‘protein kinase binding’ (**Figure 2D, bottom**). Kinases identified by examining our list of up- or down-regulated phosphosites for peptides that mapped to a kinase included members of the CaMK and MAPK families, as well as the AMPK complex.

Western blot analysis of selected synaptic and cytoskeletal proteins confirmed that the observed phosphoproteomic changes did not reflect major alterations in protein abundance, consistent with the global proteomics results (**Supplementary Figure 3**). Nevertheless, we reanalyzed the global proteomic dataset using a relaxed statistical approach, where proteins with p<0.05 (despite a q-value > 0.1) were subjected to KEGG pathway analysis. The results revealed ‘Parkinson’s disease’ as the most significantly enriched pathway, with 3 of 8 proteins within the pathway consisting of 20S proteasomal subunits (**Supplementary Figure 4A**). However, biochemical assays did not show significant differences in proteasomal activity between cortical lysates from PFF-versus monomer-injected animals (**Supplementary Figure 4B**), consistent with the non-significant abundance changes observed with the proteasomal hits (q>0.1).

### PFF-induced phosphoproteomic changes observed in the rat sensorimotor cortex are also evident in a mouse synucleinopathy model

To independently validate the PFF-mediated phosphoproteomic changes observed in the rat sensorimotor cortex, we repeated the experiment using a mouse aSyn PFF model. C57BL/6 mice were unilaterally injected with aSyn PFFs (a different preparation from the one used for the rat study) or control monomer, either in the striatum or directly in the sensorimotor cortex. After a 3-month incubation period, homogenates were prepared from the sensorimotor cortex of mice in all four groups and processed for phosphoproteomic analysis (**Figure 3A,B**). We identified approximately 3,500 phosphosites, with a larger proportion showing a trend towards up-regulation compared to down-regulation in cortical homogenates of mice injected with PFFs in the striatum or cortex (**Figure 3C,D; Supplementary Figure 5A,B**). A total of 526 or 296 sites exhibited significant changes in abundance (q-value<0.1) in the brains of mice injected with PFFs versus monomer in the striatum or cortex, respectively. As observed in the phosphoproteomic dataset derived from analyses of rat cortex, only a subset of phosphosites within individual phosphoproteins exhibited PFF-dependent dysregulation. PCA (**Figure 3E,F**) and clustered heatmap analysis (**Supplementary Figure 5C,D**) revealed a clear separation between the PFF- and monomer-treated groups, independent of the injection site, providing strong evidence of differences in the phosphoproteome profiles between PFF- and monomer-injected animals. Moreover, the phosphosite hits observed in the brains of mice injected with aSyn PFFs in the striatum or cortex were highly correlated (**Supplementary Figure 6A**), further suggesting that the observed changes are due to seeded aSyn aggregation, irrespective of the site of inoculation. These findings underscore the robustness of the observed differences and suggest that seeded aSyn aggregation induces cell signaling perturbations leading to altered phosphorylation events in the mouse sensorimotor cortex.

**Figure 3:**
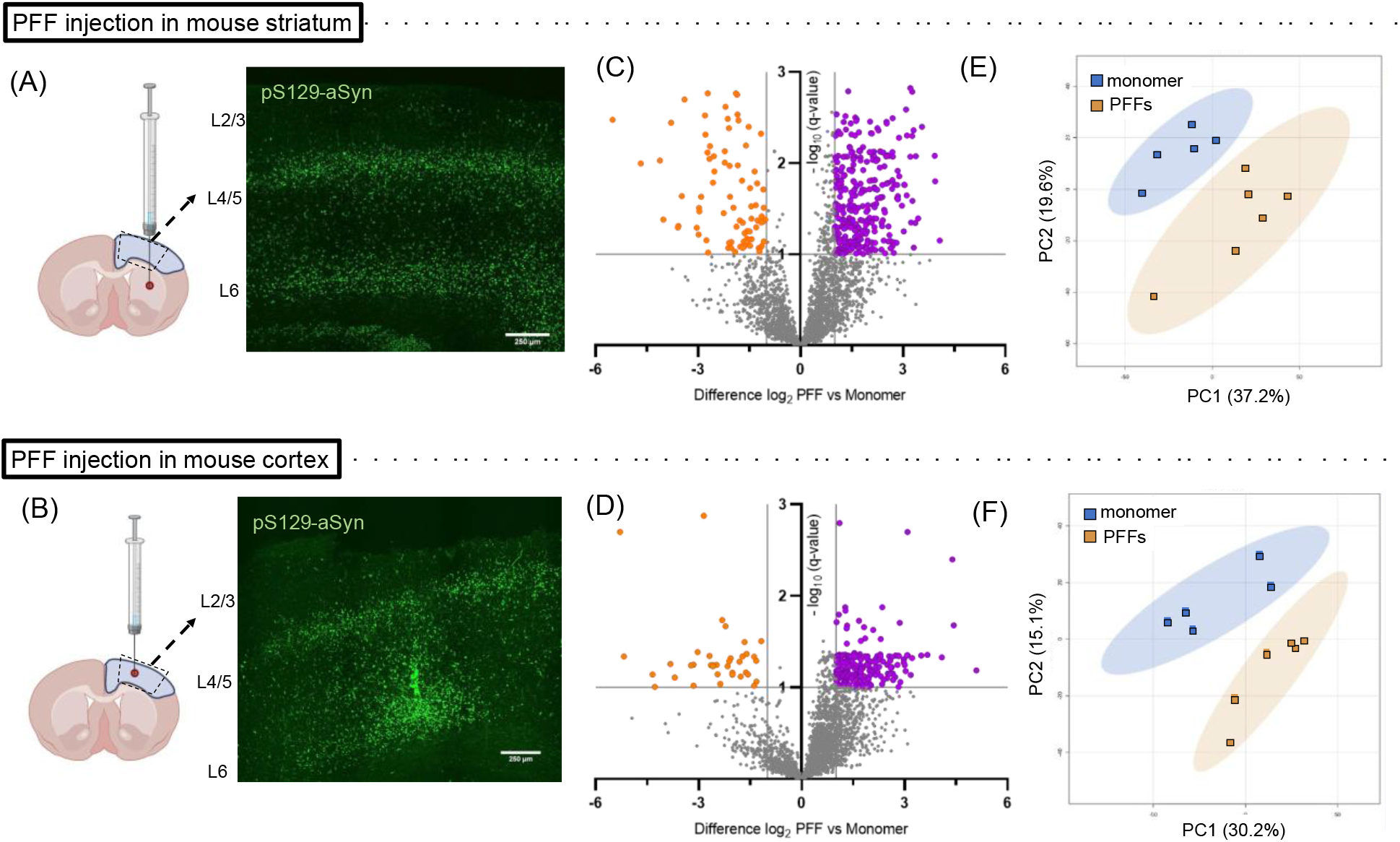
Mice injected in the striatum or cortex with aSyn PFFs show evidence of phosphoproteomic changes in the sensorimotor cortex. (A, B) Schematics showing the injection site, tissue isolated for phosphoproteomic analysis (blue-shaded region), and approximate cortical section area processed for histology (dashed trapezoid) (LEFT), and corresponding images of pS129-aSyn immunoreactivity (EP1536Y, green) (RIGHT), for mice injected with aSyn PFFs or monomer in the striatum (A) or cortex (B). The sections were prepared 3 months after injection (scale bar: 250 µm) (n = 3 animals in each group). Schematic created using https://BioRender.com. (C, D) Volcano plot distributions of phosphosites found to be up-regulated (purple) or down-regulated (orange) in homogenates prepared from mouse sensorimotor cortex 3 months after injection with aSyn PFFs versus monomer in the striatum (C) or cortex (D). The plot shows the fold change of the phosphosites in relation to the adjusted p-value (i.e. q-value), where q<0.1 was considered significant, as indicated by the horizontal dashed line. (E, F) Graphs showing the results of unassigned/unsupervised PCA of the log_2_-transformed intensities of phosphosite hits in cortical homogenates from mice injected with aSyn PFFs or monomer in the striatum (E) or cortex (F), as described in (C) and (D).

### PFF-induced aSyn aggregation leads to synaptic dysfunction and neurocircuitry perturbations in the mouse sensorimotor cortex

Although aSyn PFF models have been used extensively to study seeded aggregation in neurons, PFFs can also trigger reactive changes in glial cells (Trudler, Sanz-Blasco et al. 2021, Massari, Dues et al. 2024, Stoll, Kemp et al. 2024). Consistent with this idea, our global proteomics analysis revealed modest but significant upregulation of GFAP in the cortex of PFF-injected rats (**Supplementary Figure 7A**), which we further confirmed by immunohistochemistry (IHC) in PFF-injected mice (**Supplementary Figure 7B**). Given this evidence of glial activation, we sought to determine whether the phosphoprotein hits identified in our phosphoproteomic analysis were primarily of neuronal or glial origin. To address this question, we analyzed the cell-type specificity of the transcript-level abundance of proteins carrying significantly altered phosphosites using published single-cell transcriptomic data (Yao, van Velthoven et al. 2023). This analysis indicated that the majority of the proteins, regardless of q-value order, are neuron-enriched **(Supplementary Figure 8).** Only a modest number of hits were predominantly expressed in astrocytes, microglia, or oligodendrocytes, suggesting that the cellular origin of most of the phosphoproteomic changes is predominantly neuronal.

Next, genes associated with significantly up- and down-regulated phosphosites in cortical homogenates of PFF-treated mice (independent of injection site) were investigated via GO enrichment analysis for pathway-level information. Examination of enriched cellular localization terms revealed categories related to synaptic function (e.g., ‘post-synapse’, ‘pre-synapse’, ‘axon’, ‘dendrite’), cytoskeletal organization (e.g., ‘microtubule’, ‘structural constituent of cytoskeleton’), and protein kinase binding (**Figure 4A**). Several phosphopeptides were found to originate from the CaMK, MAPK, and AMPK families (as observed in the rat phosphoproteomics dataset), as well as the PKA family. Additionally, GO analysis of dysregulated phosphosites derived from CAMK2G, PRKAB, PRKAR, and the MAPK kinase family in our mouse phosphoproteomic datasets revealed ‘post-NMDA receptor activation events’ as the primary disrupted pathway. Similar results were obtained when the GO analysis was performed on genes associated with significantly up- and down-regulated phosphosites in cortical homogenates of mice injected in either the striatum or cortex (**Supplementary Figure 6B,C**). We also performed a protein-protein interaction (PPI) network analysis, yielding results that helped visualize potential interactions among significantly altered phosphoproteins. The resulting network map suggested that many of the observed phosphoprotein changes were interrelated and likely influenced each other (**Supplementary Figure 9**). Of note, a key altered network consisted of proteins associated with glutamatergic synapses, including Piccolo, Bassoon, Sv2A, Grin2b, Dlg2, lqSeq2, and synapsin-3 (Syn3), all of which were detected in both the striatal and cortical datasets (**Supplementary Figure 9B**). These findings are consistent with previous reports showing that aSyn aggregation occurs predominantly in excitatory neurons (Stoyka, Arrant et al. 2020). An additional altered network consisted of non-glutamatergic synaptic proteins, including synapsin-1 (Syn1). Collectively, the identified GO terms and results of the PPI network analysis further support the idea that the significant phosphoproteomic changes have a predominantly neuronal origin.

**Figure 4:**
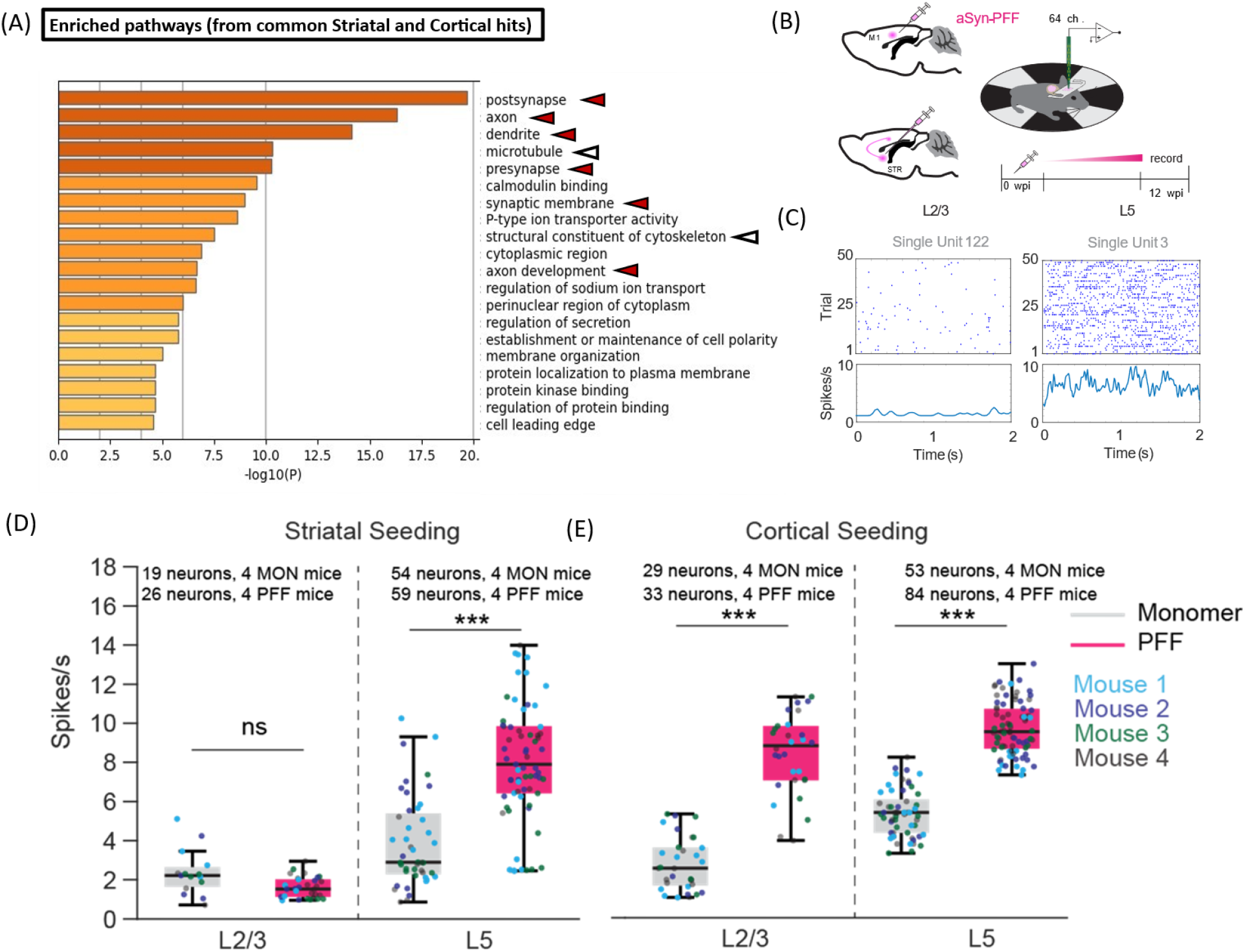
Mice injected with aSyn PFFs show evidence of cortical circuit dysfunction. (A) Chart showing enriched GO terms in the ‘cellular localization’ category that represent groups of phosphoproteins in mouse sensorimotor cortex impacted by PFF administration in either the striatum or cortex, 3 months after injection. GO analysis was carried out on proteins containing phosphosites with q<0.1 and a fold change of ≥2 in at least one group (Supplementary Figure 4A,B). Arrowheads highlight items relevant to synaptic transmission (red) or the cytoskeleton (white). (B) Schematic illustrating the design of experiments aimed at monitoring neurocircuitry function in mouse sensorimotor cortex. Two separate cohorts of mice were injected with aSyn PFFs or monomer in the primary motor cortex or dorsal striatum. Electrophysiological analyses were carried out at 0 weeks and 3 months (12 weeks) post-injection (0 and 12 wpi). (C) Example of layer-specific unit activity during awake recording in head-fixed mice injected with aSyn monomer. (D, E) Spiking rates of recorded neurons were analyzed based on cortical depth in mice injected with either aSyn PFFs or monomers into the striatum (D) or cortex (E). Number of neurons recorded per mouse injected in the striatum (D): L2/3 neurons, 3-6 (monomer) and 3-8 (PFF); L5 neurons, 8-15 (monomer) and 10-19 (PFF). Number of neurons recorded per mouse injected in the cortex (E): L2/3 neurons, 5-9 (monomer) and 6-8 (PFF); L5 neurons, 9-16 (monomer) and 11-30 (PFF). Each subgroup includes data from 4 animals. The box plot shows the median and interquartile range (IQR); *** p<0.0001, ranked-sum Wilcoxon test.

Based on these findings, we measured circuit activity as a proxy for synaptic dysfunction in the sensorimotor cortex of mice injected with aSyn PFFs or monomer. To assess whether seeded aSyn aggregation induces changes in cortical activity, we carried out silicon probe recordings in mouse primary motor cortex 3 months after PFF or monomer injection (**Figure 4B**). Quantification of spiking activity across the recording session revealed sparser firing of putative layer 2/3 (L2/3) pyramidal neurons compared to layer 5 (L5) neurons, independent of PFF or monomer administration **(Figure 4C**). In addition, we observed notable differences in firing activity across distinct layers in relation to the injection site (**Figure 4D,E**). Mice injected with PFFs in the striatum showed an increase in average spiking rates in L5 (but not L2/3) pyramidal neurons compared to monomer-injected animals (**Figure 4D**). In contrast, both L2/3 and L5 pyramidal neurons exhibited increased spiking rates in mice injected in the cortex with PFFs versus monomer (**Figure 4E**). Given that striatal aSyn PFF injection leads to pathology predominantly in layer 4/5, whereas PFF injection directly into the cortex induces extensive pathology across additional layers including L2/3 (Khan, Dutta et al. 2024), these data suggest that the observed layer-specific differences in firing rates between intrastriatally versus intracortically injected mice reflect the different patterns of aSyn propagation-induced dysfunction across the two sets of animals. Moreover, we infer that the precise location of aSyn seeding is a key determinant of the extent of hyperexcitability induced within the primary motor cortex. Neuronal hyperexcitability was also observed in the cortex of GCaMP6 mice over a period of 2 to 3 months after intrastriatal or intracortical PFF injection, as demonstrated through live calcium imaging (Khan, Dutta et al. 2024).

Next, we examined whether the observed increase in spiking activity among individual neurons in the cortex of PFF-treated mice was coordinated within a functionally connected neural network. To address this question, we computed the coherence between the spikes of any given pair of neurons within L2/3 and L5 (**Supplementary Figure 10**), yielding a frequency-resolved measure of spike coupling across these pairs. The spike-spike coherence was markedly increased across frequencies of 5 to 20 Hz in mouse cortex 3 months after intrastriatal or intracortical injection with aSyn PFFs versus control monomer (**Supplementary Figure 10B**). Taken together, these data suggest that PFF injection leads to a dynamic network transition, characterized by increased fluctuations across slow (200 ms) to intermediate (50 ms) timescales, signifying enhanced correlation within the cortical neuronal population. Thus, aSyn pathology has a complex impact on translaminar cortical activity, affecting both firing rates and network connectivity. These changes in neurocircuitry function correlate with, and could potentially be a consequence of, the perturbations of synaptic activity suggested by the phosphoproteomic data.

Because our GO analysis revealed enriched cellular localization terms related to cytoskeletal organization (**Figure 4A**), we analyzed the spine density of cortical neurites in the brains of mice injected with aSyn PFFs or monomer. Our rationale for these measurements was based on two factors: (i) cytoskeletal destabilization is closely related to a loss of neurite integrity (Dent, Merriam et al. 2011), and (ii) a decrease in dendritic spine density was previously observed in mouse cortex 5 months after intrastriatal aSyn PFF injection (Blumenstock, Rodrigues et al. 2017). Contrary to these earlier findings, we observed no significant differences in spine density between mice injected with aSyn PFFs or monomer, regardless of the injection site (**Supplementary Figure 11**). This result indicates that a decrease in spine density does not occur in our mouse aSyn PFF model at the 3-month timepoint, despite evidence of phosphoproteomic perturbations leading to altered cytoskeletal organization in the brains of these animals.

### PFF-mediated phosphoproteomic changes are predicted to involve alterations in the activity of several kinases

To gain insight into the kinases – and, therefore, the potential signaling pathways – involved in aSyn PFF-mediated phosphoproteomic alterations, we examined phosphosites that were significantly up- or down-regulated in the brains of PFF-treated rats or mice for evidence of phosphorylation motif enrichment using a sequence motif discovery approach. The analysis was carried out separately for the sequence window of phosphosites showing significant increases or decreases in phosphorylation, as determined by comparing data obtained for PFF-versus monomer-treated animals within each cohort (intrastriatally injected rats, intrastriatally injected mice, and intracortically injected mice). The results are shown in **Figure 5A**, where the frequency of individual amino acids at each position in the phosphorylation site is represented by relative size in a sequence logo. Many of the motifs that were up- or down-regulated in the brains of PFF-treated animals showed an enrichment of amino acids with negatively charged side chains, such as aspartate (D) or glutamate (E). This finding implies that acidophilic kinases, which have a high propensity to bind acidic residues in their target substrates through electrostatic interactions, exhibit altered activities in the brains of PFF-injected rats or mice.

**Figure 5:**
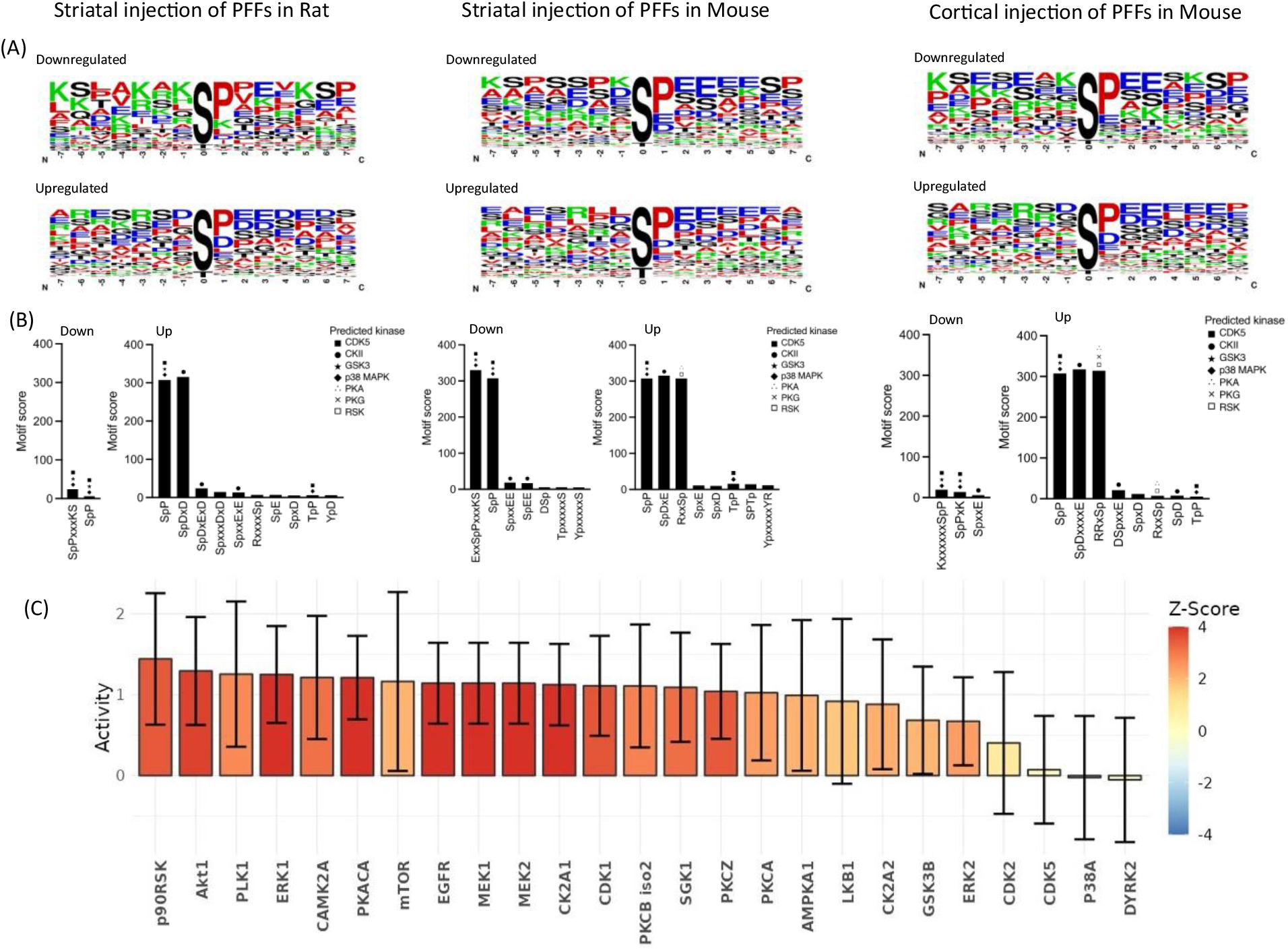
Motif enrichment and kinase prediction analyses implicate altered kinase activity in aSyn PFF-induced phosphoproteomic changes. A. Sequence logos illustrating the frequency of individual amino acids (with larger size indicating higher frequency) in the sequence windows of phosphosite-containing peptides that were down-regulated (top) or up-regulated (bottom) in the brains of rats injected with aSyn PFFs in the striatum (left) and mice injected with PFFs in the striatum (middle) or cortex (right). The phosphorylated residue is indicated at position 0. B. Bar graphs depicting enrichment scores for phosphosite motifs identified in significantly down- or up-regulated phosphosites (‘down’ or ‘up’) in the brains of rats injected with PFFs in the striatum (left) and mice injected with PFFs in the striatum (middle) or cortex (right). Higher enrichment scores indicate more significant and specific motifs. All displayed motif enrichments and kinase predictions are statistically significant (p<1 x 10^-5^ or prediction score >0.5, respectively). Symbols for predicted kinases are positioned above each bar in order of their prediction scores, with the highest score indicated by the top symbol. C. Graph showing inferred kinase activity scores generated by RoKAI analysis using phosphosites significantly altered between cortical homogenates of PFF-versus monomer-treated rats and mice (q < 0.1) and restricted to experimentally validated kinase-substrate relationships. Kinase activity scores were calculated from the coordinated phosphorylation changes of known substrates, with positive values indicating increased inferred kinase activity in PFF-injected as compared to monomer-injected rodents. Only kinases meeting the analysis thresholds of ≥3 mapped substrates and a minimum |Z|-score of 0.5 were included.

Next, we performed kinase predictions on the up- or down-regulated phosphosites within each cohort by comparing them to known consensus motifs targeted by specific kinases. Motifs that were found to be enriched in our datasets are shown in a bar graph **(Figure 5B**), with each bar manually annotated with the corresponding predicted kinases based on their prediction scores. We noted the presence of SpP (including SpP-containing) and TpP motifs with high motif scores (i.e., high levels of enrichment). However, because these proline-directed motifs tend to be overrepresented across phosphoproteomic datasets and cannot reliably distinguish between specific proline-directed kinases (e.g., GSK3, CDK5, and p38 MAPK) (Mayya and Han 2009), they provided limited mechanistic insight.

Among the other high scoring motifs, especially from up-regulated phosphosites in the brains of PFF-injected rats or mice, motifs predicted to be substrates of casein kinase 2 (CK2) were consistently enriched (e.g., SpDxD, SpDxE, and SpDxxxxE). Additionally, we identified the canonical PKA sites RxxSp and RRxSp among the list of up-regulated phosphosites in the brains of mice injected intrastriatally or intracortically with PFFs. These results suggest that aSyn pathology leads to increased activity of CK2 and PKA in rodent brain.

To support our motif-based predictions, we performed kinase activity inference (RoKAI) analysis using significantly altered phosphosites with only experimentally validated kinase– substrate relationships. Across all the unique phosphosites from the three combined datasets, the analysis identified only 121 phosphorylation sites with experimentally annotated upstream kinases (**Figure 5C**), some of which are targeted by multiple kinases. The results revealed increased activity of CK2A1 and PKACA, consistent with motif-based predictions. In addition, AKT1, known to function downstream of CK2A1, was predicted to be activated. Furthermore, the analysis revealed robust activation of the ERK signaling cascade, including the core kinases ERK1/2, their upstream activators MEK1/2, and the downstream effector p90RSK.

### PFF-induced aSyn aggregation leads to neuron-specific activation of the MAPK (ERK) signaling pathway and the formation of pERK1/2^+^ granulovacuolar body-like structures

Motivated by the RoKAI analysis results suggesting ERK pathway activation, we analyzed our list of significantly altered phosphopeptides for sequences of kinases in this pathway, focusing on known activating phosphorylation sites. Peptides phosphorylated at the canonical ERK1/2 activation sites (T202/Y204 and T183/Y185) were enriched across all three datasets (**Figure 6A**). This observation, along with the presence of multiple experimentally validated pERK1/2 substrates in our datasets (e.g., Kcnq2, PGK1), provided further evidence that the ERK pathway was activated in the cortex of PFF-injected animals. To validate these findings, Western blot analysis was performed on an independent cohort of animals (**Supplementary Figure 12A,B**). Densitometric analysis revealed a trend toward increased pERK1/2 levels in the cortex of PFF-injected mice, consistent with the phosphoproteomics data. However, this increase did not reach statistical significance, potentially due to variability in aSyn pathology burden across PFF-injected animals. To examine pERK signaling in a complementary cellular model, we used primary cortical cultures prepared from embryonic rat brains. These cultures have been shown by our group and others to form intraneuronal, Lewy-like aggregates in response to aSyn PFF treatment, recapitulating key pathological features observed in the brain (Volpicelli-Daley, Luk et al. 2011, Ivey, Guzman Sosa et al. 2025). Western blot analysis using this model demonstrated a significant increase in pERK1 levels upon PFF exposure (**Supplementary Figure 12C,D**), consistent with the activation of ERK1 inferred from our phosphoproteomic data (**Figure 6A**).

**Figure 6:**
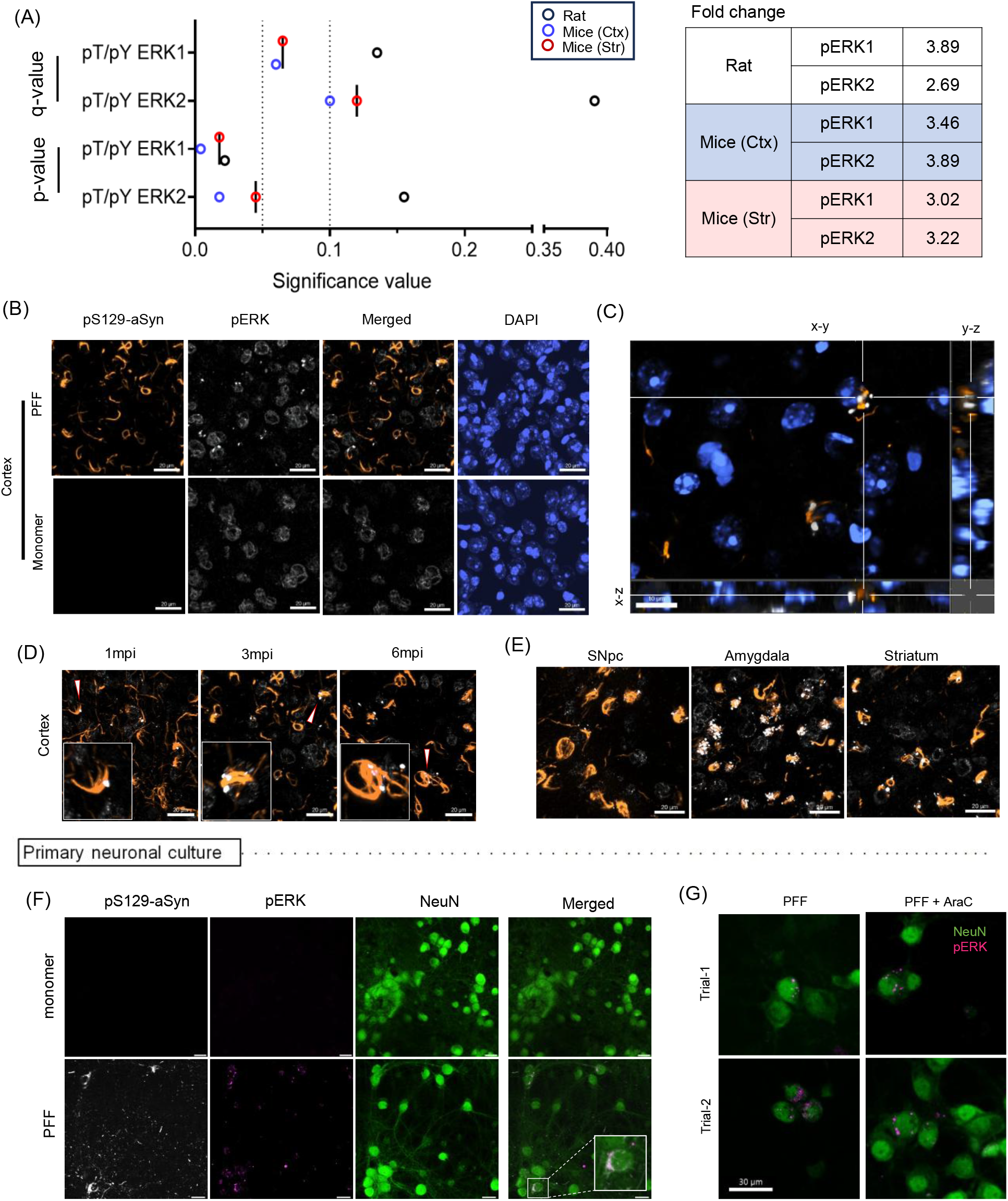
The ERK pathway is activated in aSyn aggregate-containing neurons in the brains of rodents injected intrastriatally with aSyn PFFs. A. Mean p-values and q-values (left) and fold-change values (right) for phosphorylation sites pT202/pY204 in MAPK3 (ERK1)-derived peptides and pT183/pY185 in MAPK1 (ERK2)-derived peptides, identified via phosphoproteomic analysis of sensorimotor cortex from rats injected in the striatum and mice injected in the cortex (‘Ctx’) or striatum (‘Str’) with aSyn PFFs versus monomer. B. Representative images showing increased pERK1/2 puncta in aggregate-containing cortical neurons in mice injected with aSyn PFFs versus monomer (n = 4), 3 months post-injection. C. High magnification z-stack image of DAPI, pERK1/2, and pS129-aSyn, with x-y, y-z, and x-z optical cross-sections indicating extranuclear localization of pERK1/2 in proximity to pS129-aSyn. D. Time-course analysis of pERK1/2 accumulation in aSyn aggregate-containing cortical neurons in PFF-injected mice at 1, 3, and 6 months post-injection (mpi); n = 3 or 4 animals/timepoint. Each inset shows a zoomed-in view of the neuron marked in the lower-magnification image by a white arrowhead. E. Representative images showing pERK1/2 puncta in aggregate-containing neurons in the striatum, amygdala, and SNpc of mice injected with aSyn PFFs, 3 months post-injection. F. Representative images showing increased pERK1/2 puncta in mouse cortical cultures treated with aSyn PFFs versus monomer, as well as enrichment of pERK1/2 puncta in NeuN^+^ neurons containing pS129-aSyn aggregates (n = 4). Scale bars: 25 µm. G. Representative images showing pERK1/2 puncta in AraC-treated (astrocyte-depleted) cortical cultures (N=2)

To confirm these findings and evaluate the spatial relationship between pERK1/2 and aSyn aggregate-bearing neurons, which represent only a small fraction of the total cortical neuron population, we performed IHC analyses of cortical sections from PFF-versus monomer-injected rodents. In addition to the expected nuclear pERK1/2 signal, we observed prominent granular extranuclear pERK1/2 immunoreactivity selectively in cortical neurons containing aSyn aggregates in both mice and rats injected intrastriatally with PFFs (but not monomer) at 3 months post-injection **(Figure 6B,C; Supplementary Figure 13)**. In contrast, total ERK1/2 levels remained unchanged **(Supplementary Figure 14)**, indicating that the increase in pERK1/2 levels in the brains of PFF-injected rodents arises from altered phosphorylation dynamics rather than changes in protein abundance. Collectively, these observations suggest that the PFF-dependent increase in pERK1/2 peptide abundance detected by phosphoproteomics could result from two interrelated processes: (i) the accumulation of extranuclear pERK1/2 puncta in aSyn aggregate-bearing neurons, and (ii) the activation of canonical nuclear ERK signaling

Further analyses revealed that pERK1/2⁺ neuronal puncta are present in aggregate-containing neurons across (i) multiple timepoints (1, 3, and 6 months post-injection; **Figure 6D**); and (ii) additional brain regions beyond the cortex, including the SNpc, amygdala, and striatum, in PFF-injected mice (**Figure 6E**). Similar pERK1/2 puncta were also observed in primary cortical (neuron-glia) cultures as early as 6 days post-PFF treatment, again specifically in aSyn aggregate-bearing neurons **(Figure 6F)**. This result led us to investigate the potential role of astrocytes in puncta formation. We found that prominent pERK1/2 puncta and pS129-aSyn aggregates persisted even when astrocytic content was reduced by AraC treatment **(Figure 6G**, **Supplementary Figure 15).** Collectively, these observations demonstrate that the PFF-induced formation of pERK1/2 puncta is a robust and broadly distributed phenotype that emerges rapidly even in simplified neuronal cultures, in turn suggesting that it represents an intrinsic neuronal response to aSyn aggregation.

Previous analyses of PD patient brain tissue using immunogold labeling revealed a close spatial relationship between pERK1/2 and pathological aSyn inclusions. Consistent with this observation, we found that the pERK1/2 puncta in the brains of PFF-treated animals exhibited strong colocalization with pS129-aSyn aggregates in neurons **(Figure 6B-E)**. The morphology and predominantly extranuclear distribution of these puncta were reminiscent of granulovacuolar bodies (GVBs), dense protein-rich vacuolar structures associated with cellular stress and impaired lysosomal autophagy. GVBs have been characterized primarily in proximity to fibrillar tau tangles in pyramidal neurons of Alzheimer’s disease (AD) and tauopathy models. To determine whether the pERK1/2 puncta observed in our aSyn PFF models correspond to GVB-like structures, we stained cortical sections from PFF-injected animals for various proteins previously associated with GVBs. We observed robust colocalization of pERK1/2 puncta with the canonical GVB markers CK1δ and CHMP2B, as well as S202/pT205 Tau **(Figure 7A)**. Importantly, all three markers were enriched in aggregate-bearing neurons and showed a similar close spatial association with pS129-aSyn (**Figure 7B**) as reported for GVBs near tau aggregates in AD pyramidal neurons. The association between pERK1/2 and GVB-like structures was further validated in cultured neurons exposed to aSyn PFFs (**Figure 7C**), demonstrating that this phenomenon is reproducible across experimental systems. Together, these findings establish extranuclear pERK1/2 as a reliable marker of GVB-like structures associated with aSyn pathology, thereby linking ERK1/2 signaling to an early neuronal stress response in the context of aSyn aggregation.

**Figure 7.**
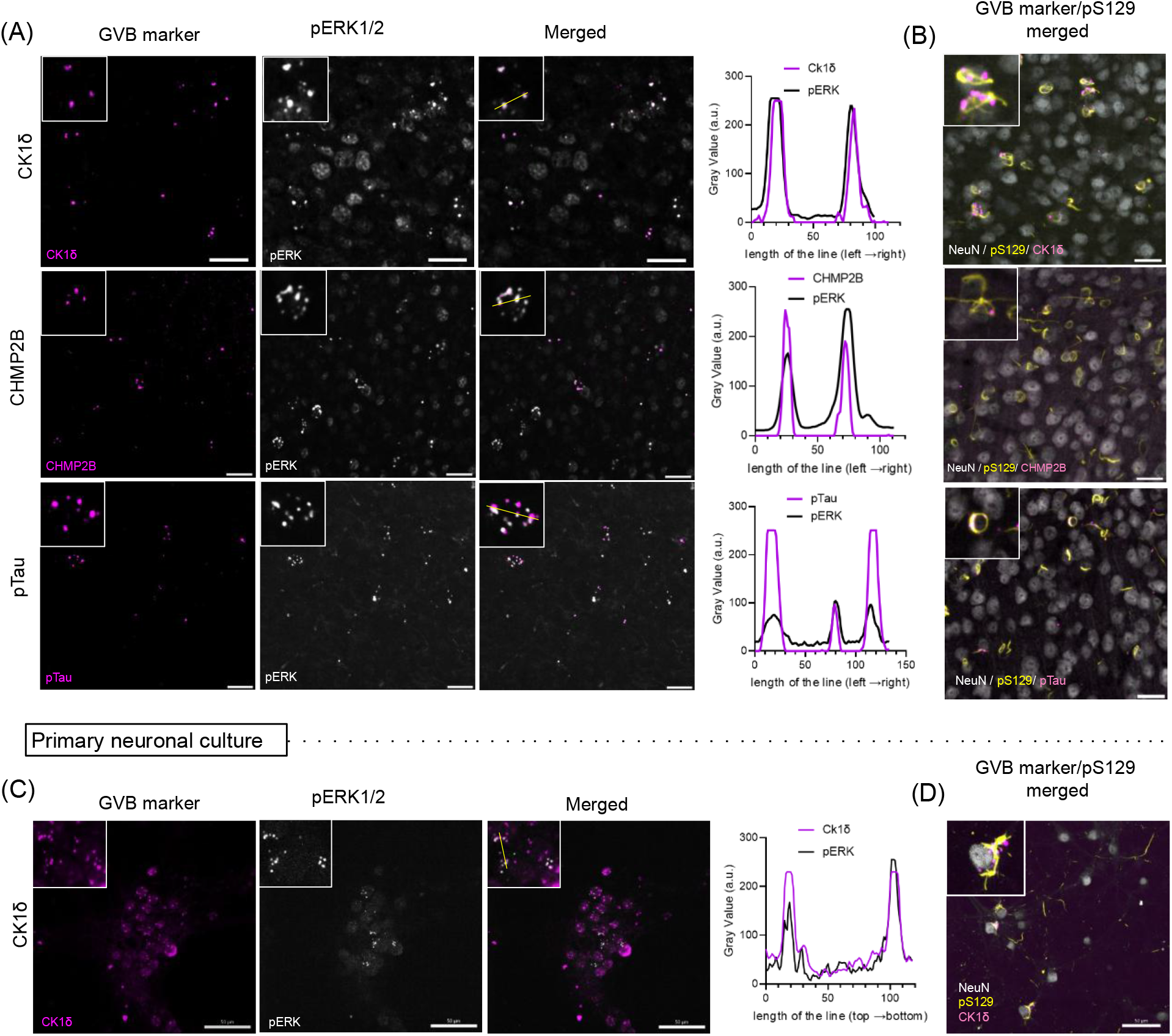
Extranuclear pERK1/2 puncta exhibit features consistent with granulovacuolar body (GVB)-like structures in aSyn aggregate-bearing neurons. A. Confocal images of cortical sections from PFF-injected mice showing co-localization of pERK1/2 (white) with the canonical GVB markers CK1δ and CHMP2B, or pTau (pS202/pT205) (magenta). Merged images reveal overlap of pERK1/2 puncta with these markers. Right: line intensity profiles (yellow line) illustrating spatial co-distribution. B. Images of cortical sections from PFF-injected mice showing association of GVB markers with pathological aSyn. Neurons (NeuN^+^, gray) exhibit co-localization of pS129-aSyn (yellow) with CK1δ, CHMP2B, or pTau (pS202/pT205) (magenta), indicating that GVB-like structures are formed in aSyn aggregate-bearing neurons. C. Images of PFF-treated cortical cultures showing co-localization of pERK1/2 (white) with CK1δ (magenta). Right: line intensity profiles (yellow line) showing spatial overlap. D. Image of PFF-treated cortical cultures showing co-localization of CK1δ with pathological aSyn. Neurons (NeuN^+^, gray) exhibit overlap of pS129-aSyn (yellow) with CK1δ (magenta), confirming formation of GVB-like structures in aggregate-bearing neurons *in vitro*. Insets in panels (A)-(D) show higher-magnification views of individual puncta. Scale bars: 25 μm (A, B); 50 μm (C, D).

## Discussion

The spread of aSyn pathology in the brains of rodents injected intrastriatally with PFFs follows a pattern determined by the anatomical connectivity of different brain regions to the striatum (Henderson, Cornblath et al. 2019). Of relevance to the current study, PFFs injected in the striatum are taken up by corticostriatal neurons via endocytosis in the synaptic region. Once internalized, PFFs are thought to escape from the endocytic compartment and induce the aggregation of cytosolic aSyn, which is abundant at the synaptic terminal of the recipient neuron (Burre 2015). Aggregates then spread gradually in a retrograde manner from the synapse to the cell body. This spread of aSyn pathology is cell type-specific, with excitatory neurons of layer IV/V (**Figure 1C**) being particularly susceptible to aggregate formation and propagation (Bopp, Holler-Rickauer et al. 2017, Stoyka, Arrant et al. 2020, Goralski, Meyerdirk et al. 2024), closely mimicking the pathology observed in human PD (Harding and Halliday 2001).

Although cortical pathology becomes evident in rodents within 15 days of intrastriatal PFF administration (Khan, Dutta et al. 2024), motor and cognitive deficits appear much later (Luk, Kehm et al. 2012, Patterson, Duffy et al. 2019, Stoyka, Arrant et al. 2020s). This slow disease progression and the modest initial behavioral phenotype closely mirror the early stages of synucleinopathy diseases such as PDD and DLB (Schulz-Schaeffer 2010). Accordingly, the rodent aSyn PFF model is well-suited for investigating early cortical cellular events that precede overt neurodegeneration. To this end, we carried out proteomic, phosphoproteomic, and lipid profiling analyses of brain homogenates from rats and mice injected intrastriatally or intracortically with aSyn PFFs, 3 months post-injection. Unexpectedly, the global proteomic analyses revealed only a small number of proteins with significant changes in abundance in the brains of PFF-injected animals across various brain regions, based on p-values but not q-values from FDR-corrected significance tests (**Figure 1E**; **Supplementary Figure 1F, G**). Comparing our results with data from one previous global proteomic analysis of aSyn PFF-injected mice revealed substantial overlap between our list of dysregulated proteins and those identified in the earlier study (Ma, Seo et al. 2021). The limited global proteomic differences observed between PFF- and monomer-injected rats may reflect insufficient statistical power, given the high variability of the global proteomics dataset, or they may indicate that widespread expression changes emerge only at later time points post-injection. An additional possibility is that our global proteomic analyses were performed on cortical tissue from PFF-injected rats rather than mice. Notably, rats showed a lower pathology burden than mice (**Figure 1C vs. Figure 3A,B**), likely due to their 3 to 4-fold greater brain volume, which was not fully offset by a 2-fold increase in PFF dose (see ‘Experimental Methods’). This lower pathological burden likely translates into a reduced effect size in rats, which may not be overcome simply by increasing animal numbers (i.e., study power). For instance, GFAP was significantly altered (p < 0.017) in the global rat dataset but did not cross the typical >2-fold cutoff (i.e., only 1.62-fold upregulation). If the fold-change criterion is relaxed, 5.4% of proteins (116 total) with p < 0.05 map to PD-relevant pathways (**Supplementary Figure 4**) and overlap with prior reports. This issue with statistical power and lower effect size may have also contributed to the lack of significantly dysregulated lipids revealed by our lipid profiling analysis of brain homogenates from PFF-injected rats (**Figure 1I, Supplementary Figure 2**).

In contrast to the low hit rate observed in our global proteomic study, phosphoproteomic analysis led to the identification of 220 phosphosites from 161 proteins that were differentially regulated in cortical homogenates of PFF-versus monomer-injected rats, with most being up-regulated ≥ 2-fold (**Figure 2A**). Similar results were obtained with mice injected with aSyn PFFs in the striatum or cortex (**Figure 3; Supplementary Figure 5**). This overlap in cortical phosphoproteomic profiles between striatally and cortically seeded mice indicates that, by 3 months post-injection, both paradigms produce a cortex enriched with intracellular aSyn aggregates and associated signaling perturbations – regardless of whether the pathology reached cortical neurons via long-distance retrograde spread from the striatum or through direct local seeding. This convergence strongly supports the view that the observed phosphoproteomic changes are driven primarily by the presence of aggregated aSyn within neurons themselves, rather than by the specific route of propagation. Among the differentially phosphorylated proteins observed in PFF-treated rats or mice were pre- and post-synaptic scaffolding proteins, proteins involved in synaptic vesicle pool regulation and release, and proteins involved in postsynaptic signaling. GO analysis of the proteins associated with the top differentially regulated phosphosites further validated disruptions of synaptic function in the brains of PFF-versus monomer-injected animals (**Figures 2D, 4A; Supplementary Figure 6B,C**). Changes in cellular pathways involved in synaptic function have also been observed in global transcriptomics and proteomics analysis of brain samples from aSyn PFF-injected mice, transgenic PD mice, and PD patients (Bridi and Hirth 2018, Goralski, Meyerdirk et al. 2024, Patterson, Kochmanski et al. 2024, Gao, Chlebowicz et al. 2025). However, in contrast to these studies, we failed to observe changes in total synaptic protein expression levels in rats (**Supplementary Figure 3**). This discrepancy may reflect a divergence between mRNA and protein expression changes, as reported by others (Mangleburg, Wu et al. 2020, Tasaki, Xu et al. 2022).

Previous studies have demonstrated that acute synaptic depolarization significantly alters the synaptic phosphoproteome (Kohansal-Nodehi, Chua et al. 2016, Engholm-Keller, Waardenberg et al. 2019). Here, we show that chronic protein aggregation similarly disrupts the phosphorylation states of synaptic proteins. However, we observed minimal overlap between the phosphosites altered in our study and those reported from earlier analyses of acute depolarization. In addition, the serine phosphorylation sites identified here were not significantly enriched for the RxxS motif, a known CaMKII target during Ca^2+^ influx (Kohansal-Nodehi, Chua et al. 2016, Engholm-Keller, Waardenberg et al. 2019) caused by acute depolarization. Together, these results suggest that the phosphoproteomic changes observed in the brains of PFF-treated rats or mice are likely driven by chronic, PFF-induced signaling disruptions rather than acute synaptic activation. These findings are consistent with previous reports of aberrant synaptic connectivity in brain samples from PD or DLB patients, as well as in rodent synucleinopathy models, where the degree of synaptic dysregulation is associated with aggregate burden (Bridi and Hirth 2018, Froula, Henderson et al. 2018, Chen, Nagaraja et al. 2022). aSyn is known to localize to the synaptic terminal, where it interacts with other synaptic proteins and plays a key role in synaptic vesicle homeostasis (Burre 2015, Parra-Rivas, Madhivanan et al. 2023, Ramalingam, Jin et al. 2023). As one potential scenario, we suggest that the recruitment of monomeric aSyn into aggregates displaces it from its normal synaptic location (Chen, Nagaraja et al. 2022, Calabresi, Di Lazzaro et al. 2023), leading to altered synaptic protein distribution, disrupted activity, and changes in synaptic phosphorylation, ultimately impairing synaptic function and causing circuit irregularities. Similar to the results presented here, phosphoproteomic analysis of cortical samples from AD patients at various stages of disease revealed changes in cellular pathways related to synaptic function (Ferrer, Andres-Benito et al. 2021). This finding suggests that dysregulation of synaptic phosphoproteins in the cortex may be a common molecular feature underlying cognitive deficits associated with neurodegenerative disorders such as PD and AD.

Additional differentially regulated phosphopeptides in the brains of PFF-treated rats or mice were derived from proteins with molecular functions related to cytoskeletal homeostasis (**Figures 2D, 4A; Supplementary Figure 6B, C**). Among the most highly phosphorylated proteins in this category were Map1a, Map1b, and Map2, key components of pre-and post-synaptic terminals (Goralski, Meyerdirk et al. 2024). PFF-mediated perturbations of these and other cytoskeletal proteins could account for previously reported decreases in axonal transport in primary neurons treated with aSyn PFFs (Volpicelli-Daley, Luk et al. 2011).

The dysregulation of synaptic and cytoskeletal phosphoproteins in PFF-injected rodents suggested that neurocircuitry function might be altered in the brains of these animals. In support of this idea, we observed lower spike rates and altered spike coherence in the sensorimotor cortex of mice injected with aSyn PFFs, either intrastriatally or intracortically (**Figure 4D,E**). Additional studies by our group revealed that the changes in neurocircuitry function observed here at the 3-month timepoint can appear as early as 4 weeks post-PFF injection (Khan, Dutta et al. 2024) and are characterized by increases in the amplitude of ‘beta events’ (spontaneous bursts of beta transients) driven by NMDA receptor and voltage-gated Ca^2+^ channel (VGCC) signaling (Khan, Dutta et al. 2024). Our phosphoproteomic data provide a molecular basis for these neurocircuitry disruptions, implicating dysregulation of NMDA receptor-related phosphoproteins such as Grin2a/2b, Shank3, and the Dlg family in PFF-injected mice (see Figures S10 and S11 in ref. (Khan, Dutta et al. 2024)), as well as the VGCC subunits Cacna1b and Cacnb1 in PFF-injected rats (this study). These findings are also consistent with our network analysis of up- or down-regulated phosphoproteins, which revealed a cluster enriched with proteins associated with glutamatergic synapses (**Supplementary Figure 9B**). Moreover, synapsin-1 and -3, key nodes in the identified PPI network, may represent important modulators of neurocircuitry function in the mouse PFF model, as they are known to regulate the release of glutamate and other neurotransmitters. Among the synapsin-1 and -3 phosphorylation sites detected in our dataset, only a subset showed significant alterations in PFF-treated animals (**Supplementary Table 2**), suggesting that these sites may be functionally important. Future research will aim to determine whether the observed phosphorylation changes in synaptic proteins, cytoskeletal components, NMDA receptor proteins, VGCC subunits, and synapsin proteins contribute directly to the neurocircuitry alterations in the brains of seed-injected rodents. Additionally, given the cellular complexity of the sensorimotor cortex, follow-up work is needed to establish the neuronal subtype specificity of these phosphoproteomic changes.

Kinase prediction based on enriched consensus motifs as well as analysis of experimentally validated kinase-substrate pairs from the up- or down-regulated phosphosites suggested increased CK2, ERK1/2, and PKA activities, as well as a less robust decrease in CDK5 and p38 MAPK activities in the brains of PFF-injected rats or mice (**Figure 5**). Possible PFF-dependent up-regulation of CK2 activity is particularly compelling given this kinase’s association with PD pathology. Specifically, CK2 phosphorylates aSyn on residue S129, a hallmark modification of Lewy bodies and neurites in the brains of synucleinopathy patients (Okochi, Walter et al. 2000, Anderson, Walker et al. 2006). Furthermore, the CK2 β subunit has been shown to co-localize with pS129-aSyn aggregates in the SNpc (Ryu, Kim et al. 2008). As one possible mechanism, changes in CK2β phosphorylation at residues S3, S4, and S209, as observed in our phosphoproteomic dataset, could contribute to the increased CK2 activity in the brains of PFF-injected rats or mice (Litchfield, Lozeman et al. 1991, Zhang, Vilk et al. 2002).

Further analysis of the significantly altered phosphopeptides revealed kinase-derived peptides from the MAPK (ERK) family, along with the CaMK and PKA families and the AMPK complex, in the brains of PFF-injected rats or mice. Activation of the ERK pathway has not been previously demonstrated in an aSyn PFF model or in other models of inducible aSyn aggregation. The evidence presented here of increased MAPK3 (ERK1) phosphorylation on residues T202 and Y204 is consistent with previous human postmortem data showing enhanced MAPK3 phosphorylation at these sites within intraneuronal granular microstructures in the cortex of DLB patients (Fujita, Homma et al. 2018) and the SNpc of PD or DLB patients (Zhu, Kulich et al. 2002). Furthermore, our observation that punctate pERK1/2 structures are detectable as early as 1 month post PFF injection, well before significant neuronal loss, suggests that pERK1/2 accumulation is an early event linked to aSyn aggregation, consistent with data from a previous study of human PD brain samples (Zhu, Kulich et al. 2002).

Evidence that MAPK1 and other kinases such as PRKACA are associated with pS129-aSyn in Lewy bodies (Zhu, Kulich et al. 2002, Killinger, Marshall et al. 2022) implies that interactions between these kinases and pathological aSyn could play a role in the altered kinase activity driving the phosphoproteomic changes observed in the brains of PFF-treated rodents. Activation of the ERK pathway, previously linked to a loss of viability of iPSC-derived cortical neurons (Suzuki, Egawa et al. 2024), may contribute to the cortical dysfunction observed in PFF-injected mice (Figure 4B-D). Our finding that the MAPK substrates Kcnq2 and Pgk1 exhibit increased phosphorylation in cortical homogenates from PFF-treated rodents provides insights into how enhanced MAPK activity could modulate neuronal health and function in the brains of these animals. Notably, MAPK-mediated phosphorylation of Kcnq2 on residue S448, a modification that shifts the voltage sensitivity of Kcnq2/3 heteromultimeric channels to a more depolarized state, could reduce potassium channel activity, thereby increasing excitability and glutamate sensitivity(Tsuboi, Otsuka et al. 2022). Conversely, activated MAPK could enhance neuroprotection by phosphorylating Pgk1 at S203, inducing Pgk1 mitochondrial localization that, in turn, leads to reduced oxidative stress through the inhibition of mitochondrial pyruvate metabolism (Li, Jiang et al. 2016, Zhang, Sun et al. 2023). Lastly, our observation that ‘post-NMDA receptor activation events’ was the primary disrupted pathway from a GO analysis of dysregulated phosphopeptides derived from the MAPK kinase family as well as CAMK2G, PRKAB, and PRKAR suggests that these kinases could contribute to the PFF-mediated dysregulation of NMDA receptor signaling discussed earlier.

Our observation that pERK1/2 is associated with GVB-like structures in the brains of PFF-injected rodents and in PFF-treated cortical neurons provides direct evidence of cellular stress and altered proteostasis linked to aSyn aggregation, in line with observations from AD and tau PFF models (Kohler 2016, Wiersma, van Ziel et al. 2019). Notably, these structures appear as early as 1 month post-injection *in vivo* and within 6 days of PFF treatment in primary neurons, indicating that the processes driving GVB formation are activated at early stages of pathology, consistent with evidence from tau propagation studies (Wiersma, van Ziel et al. 2019, Wiersma, Hoozemans et al. 2020). Our results also demonstrate that GVB formation is neither exclusive to tau pathology (see also ref. (Dues, Erb et al. 2025)), nor restricted to pyramidal neurons, but instead reflects a conserved neuronal response to proteinopathic stress. These parallels with AD and tau PFF models support the idea that aSyn aggregates can act as a central driver of GVB formation, analogous to the role of tau aggregates (Wiersma, Hoozemans et al. 2020, Jorge-Oliva, Smits et al. 2022). Such mechanistic overlap suggests that pathways implicated in tau pathology, including upstream Ras signaling (Ferrer, Blanco et al. 2001) and downstream translational dysregulation (Castellani, Gupta et al. 2011), may be relevant to the development of aSyn pathology. Finally, the co-localization of phosphorylated αSyn and tau within the GVB-like structures observed in our study (**Figure 7A, bottom**) may help explain the frequent aSyn-tau co-pathology observed in DLB.

Although our transcriptomic comparison (**Supplementary Figure 8**) and Gene Ontology analyses (**Figures 2, 4**; **Supplementary Figure 6**) strongly support a substantial neuronal contribution to the observed phosphoproteomic changes, it remains unclear to what extent these alterations arise directly from PFF-induced aSyn aggregation within neurons versus secondary signaling mechanisms involving activated glia. Several lines of evidence suggest that intraneuronal aggregation is an important driver of these effects: (i) pERK1/2 puncta are readily detected in the amygdala (**Figure 6E**), a region showing minimal glial activation (Massari, Dues et al. 2024); and (ii) GVBs form in primary neuronal cultures largely devoid of glia following PFF-induced aSyn aggregation (**Figure 6G**), consistent with findings in tau pathology models (Wiersma, van Ziel et al. 2019). Together, these findings support a major contribution of intraneuronal aSyn aggregates to the observed phosphoproteomic changes and circuit dysfunction. However, given the cortical glial activation observed at 3 months post-injection (**Supplementary Figure 7**) and the ubiquitous expression of key kinases such as CK2 in astrocytes (Rosenberger, Morrema et al. 2016), we cannot exclude significant glial contributions to the observed phosphoproteomic signatures. Such contributions could arise both through cell-autonomous changes in glial kinase activity that directly influence the overall phosphoproteomic profile, and through alterations in glial signaling or homeostatic functions that secondarily modulate neuronal signaling pathways and phosphoproteomic states. Because aSyn aggregation and glial activation occur in parallel, their relative contributions remain difficult to fully disentangle *in vivo*. Future studies will be important for resolving the cell type-specific origins and propagation of these signaling changes.

Importantly, several of the phosphoproteomic and network-level alterations identified in our model **(Supplementary Figure 9)** are supported by data from human synucleinopathy studies, strengthening the translational relevance of our findings. In particular, Syn3 – a target of ERK1/2, PKA, and CDK5 signaling that regulates neurotransmitter release – was identified in an analysis of network alterations associated with PDD and DLB (Shantaraman, Dammer et al. 2024) and was shown to co-localize with Lewy pathology in the brains of PD patients (Longhena, Faustini et al. 2018). Syn3 has also emerged as a potential therapeutic target through its ability to modulate aSyn aggregation (Faustini, Longhena et al. 2018, Faustini, Longhena et al. 2022). More broadly, the protein networks revealed by our PPI analysis overlap with those identified via proteomic analyses of human brain tissue (Shantaraman, Dammer et al. 2024). Furthermore, the pERK1/2^+^ intraneuronal bodies identified in cortical cultures and rodent brain resemble intraneuronal granules with enhanced MAPK3 phosphorylation detected in DLB cortex (Fujita, Homma et al. 2018) and the SNpc of PD and DLB patients (Zhu, Kulich et al. 2002) and may represent stress-activated GVB-like structures similar to those associated with other neurodegenerative diseases such as AD (Kohler 2016, Wiersma, van Ziel et al. 2019). Beyond validation in human patient samples, future studies using humanized mouse models inoculated with human-derived fibrils (Yang, Shi et al. 2022, Sokratian, Zhou et al. 2024) will yield further insight into the disease relevance of these signaling and proteomic alterations.

In conclusion, our findings demonstrate that seeded aSyn aggregation induces widespread alterations in protein phosphorylation in the sensorimotor cortex of rats and mice injected with aSyn PFFs. These phosphoproteomic changes disrupt signaling pathways linked to NMDA receptor function, VGCC activity, and cytoskeletal dynamics and are accompanied by neurocircuitry dysfunction. The observed perturbations reflect the altered activity of multiple kinases, including MAPK (ERK1/2), PKA, and CK2, highlighting the complex interplay of signaling pathways underlying aSyn-mediated neuronal dysfunction. Our results further underscore the utility of phosphoproteomics for capturing molecular events associated with synucleinopathy progression, including ERK pathway activation and the emergence of GVB-like structures indicative of cellular stress, while supporting the relevance of the aSyn PFF model for recapitulating early pathogenic changes. Elucidating mechanisms linking aSyn-induced phosphoproteomic remodeling to neurocircuitry impairment may inform therapeutic strategies aimed at alleviating cognitive decline in PD and related synucleinopathies.

## Experimental Methods

### Recombinant aSyn protein purification

Mouse recombinant aSyn was purified as described (Zhang, Griggs et al. 2013, Volpicelli-Daley, Luk et al. 2014). *E. coli* BL21 (DE3) cells transformed with the bacterial expression vector pT7-7 encoding mouse aSyn were incubated in LB medium supplemented with ampicillin (100 μg/mL) at 37°C until the optical density at 600 nm reached 0.5-0.6. Protein expression was induced with isopropyl β-D-1-thiogalactopyranoside at a final concentration of 1 mM, followed by incubation at 37°C for 4 h. The cells were then harvested by centrifugation, resuspended in lysis buffer (10 mM Tris HCl, pH 8.0, 1 mM EDTA, 0.25 mg/mL lysozyme), and lysed using a French press cell disruptor (Thermo Electron, Waltham, MA) at > 1000 psi. The lysate was treated with 0.1% (w/v) streptomycin sulfate to precipitate DNA and then clarified by centrifugation. The supernatant was subjected to partial purification via two successive ammonium sulfate precipitations (30% and 50% w/v saturation) at 4°C. The resulting pellet was resuspended in 10 mM Tris-HCl (pH 7.4), and the suspension was boiled at 95°C for 15 min. Following centrifugation at 13,500 × g at 4°C for 20 min to precipitate denatured proteins, aSyn was purified from the supernatant via successive fractionations using: (i) a HiLoad 16/600 Superdex 200 pg size exclusion column (Cytiva, Marlborough, MA), with elution performed in 10 mM Tris HCl (pH 7.4); and (ii) a HiPrep Q HP 16/10 or DEAE anion exchange column (Cytiva), with elution performed using a linear gradient of 25 mM to 1 M NaCl in a buffer consisting of 10 mM Tris HCl (pH 7.4) and 1 mM EDTA. Fractions enriched with aSyn or aSyn-mVenus (identified via SDS-PAGE with Coomassie blue staining) were pooled, and the solution was dialyzed against PBS (10 mM Na_2_HPO_4_, 1.8 mM KH_2_PO_4_, 2.7 mM KCl, and 137 mM NaCl, pH 7.4). aSyn preparations used for PFF formation were depleted of endotoxin using the Pierce high-capacity endotoxin removal resin (Thermo Fisher Scientific #88277, Waltham, MA) to 0.018 units/μg of protein (3). The purified protein was stored at -80°C until use, with a final purity of approximately 95%.

### Preparation of aSyn PFFs

A 500 μL solution of monomeric mouse aSyn (5 mg/mL) was filtered through a 0.22 μm filter (Corning #8160, Corning, NY) and incubated in a sterile 1.5 mL microcentrifuge tube at 37 °C for 7 days with continuous agitation at 1,000 rpm (123 x g) in a Thermomixer (BenchTop Lab Systems #BT917, St. Louis, MO). The resulting fibrils were concentrated by centrifugation at 13,000 x g for 10 min, and the pellet was resuspended in 250 μL of Dulbecco’s phosphate-buffered saline (DPBS; Cytiva). An aliquot of the fibril suspension (5 µL) was incubated with 8 M guanidine hydrochloride at 22°C for 1 h to dissociate the fibrils into monomers. The aSyn concentration was determined via absorbance measurements at 280 nm using a Nanodrop spectrophotometer, with an extinction coefficient of 7450 M^-1^ cm^-1^. aSyn fibrils were stored at −80°C as 25 μL of aliquots at a concentration of 5 mg/mL. Prior to use, aliquots were thawed and sonicated in ethanol-sterilized sonicating tubes (Active Motif #53071, Carlsbad, CA) using a cup horn sonicator (Qsonica #q700, Newtown, CT) at 30% power (∼100 W/s) with a cycle of 3 s on and 2 s off, for a total ‘on’ time of 15 min, while maintaining the bath temperature between 5 and 15 °C.

### Transmission electron microscopy

Recombinant aSyn fibrils were analyzed by negative stain transmission electron microscopy (TEM) as described (Sanyal, Dutta et al. 2019, Landeck, Strathearn et al. 2020). Briefly, 3 μL of fibril suspension (≤ 0.5 mg/mL) was applied to a glow-discharged, carbon-coated copper TEM grid (Electron Microscopy Sciences #CF400CU, Hatfield, PA) and incubated at 22°C for 45 s. The grid was rinsed with deionized water, and 3.5 μL of 1% (w/v) phosphotungstic acid (PTA) was applied as a contrast agent. After a 1-minute incubation, excess PTA was removed by blotting with Whatman filter paper (Cytiva), and the grid was air-dried. Imaging was performed using an FEI Tecnai T12 Transmission Electron Microscope (Thermo Fisher Scientific) operating at 80 kV. Gatan DigitalMicrograph software (Gatan Microscopy Suite, Pleasanton, CA) was used to capture the images.

### Intracranial PFF injections

All methods for working with animals were conducted using protocols approved by the Purdue Animal Care and Use Committee (PACUC). In one set of experiments, 3- to 4-month-old male Sprague Dawley rats (Envigo, Indianapolis, IN) were secured in a stereotaxic frame (Kopf Instruments, Tujunga, CA) and anesthetized with isoflurane. A suspension of sonicated aSyn PFFs or a solution of control, monomeric protein (4 μL, 5 mg/mL) was injected into the striatum (coordinates: ± 3.5 mm lateral, 0 mm posterior to bregma; 4.5 mm ventral from dura) at a constant flow rate of 0.5 μL/min using a 10 μL Hamilton syringe (Hamilton, Reno, NV) fitted with a 30-gauge needle with a 45° angled tip. The needle was left in place for 5 min post-infusion to prevent backflow. In a second set of experiments, 3- to -4-month-old male C57BL/6 J mice (The Jackson Laboratory #000664, Bar Harbor, ME) underwent stereotaxic injections of aSyn PFFs or monomer (5 mg/mL) into either the primary motor cortex (1 µL delivered at ± 1.8 mm lateral, 0.3 mm anterior to bregma; 0.7 mm ventral from dura) or the striatum (1.5 µL delivered at ± 2 mm lateral, 1 mm posterior to bregma; 2.5 mm ventral from dura). Injections were performed using a 10 µL syringe fitted with a 33-gauge needle at a flow rate of 50 to 100 nL/min. Post-surgical care included administration of ketoprofen (5 mg/kg in normal saline) to rats and 5 mg/kg carprofen plus 6 mg/kg dexamethasone to mice, immediately after surgery and 48 h later.

### Brain tissue processing

Rats or mice were euthanized by an overdose with sodium pentobarbital (50.9 mg/mL) and transcardially perfused with cold PBS. Animals designated for immunohistochemical (IHC) analysis were additionally perfused with 4% (w/v) paraformaldehyde (PFA) in PBS. Brains were quickly removed after decapitation, post-fixed overnight in the same PFA solution, and cryoprotected in 30% (w/v) sucrose in water at 4°C. Coronal sections (40 μm thick) were prepared using a frozen sliding microtome (Thermo Scientific Microm HM430). Sections were stored at -20°C in cryoprotectant solution (30% w/v sucrose and 30% v/v ethylene glycol in 0.1 M phosphate buffer, pH 7.4) until IHC analysis. For animals designated for proteomic and lipid profiling, specific brain regions (sensorimotor cortex, amygdala, and SN) were isolated immediately after perfusing with cold PBS, flash-frozen in liquid nitrogen, and stored at -80°C until further analysis.

### Primary neuronal culture treatments

Primary cortical cultures were prepared from embryonic day 17 (E17) Spraque Dawley rats or C57BL/6J mice using methods approved by the IACUC, as previously described (Volpicelli-Daley, Luk et al. 2014). Isolated brain regions were dissociated using papain (Worthington Biochemical, #NC9597281), and cells were plated onto poly-L-lysine-coated culture plates at a density of 700,000 cells/well in 6-well plates (for immunoblotting) or 150,000 cells/well in 24-well plates (for immunocytochemistry (ICC)). For immunoblotting analysis, rat primary cortical neurons were incubated with 6 µg/mL aSyn PFFs for 7 days. After treatment, cells were washed with ice-cold PBS and stored at -80 °C until lysis. For ICC, cells were fixed in 4% (w/v) PFA in PBS (pH 7.4) and stored in 1× PBS at 4°C until further processing for immunostaining.

### IHC and ICC

Free-floating sections were washed 3 times for 10 min each in PBS to remove residual cryoprotectant solution and then incubated in PBS containing Triton X-100 (1% v/v) for 60 min. The sections were blocked in a solution of 10% (v/v) normal donkey serum in PBS supplemented with Triton X-100 (0.3% v/v) (PBST) for 90 min and then incubated with primary antibody solution prepared in PBST containing 1% (v/v) normal donkey serum overnight at 4°C. After washing in PBS (3 x 10 min), the sections were incubated with Alexa fluor-conjugated secondary antibodies (Jackson ImmunoResearch Laboratories, West Grove, PA; 1:500 dilution) at 22°C for 90 min. After a final round of three 10-min washes in PBS, the tissues were mounted on slides, allowed to dry overnight, and sealed with coverslips using DPX mounting media (Electron Microscopy Sciences #13512).

For ICC, fixed primary neuronal cultures were permeabilized and blocked in PBS containing 0.3% (v/v) Triton X-100, 1% (w/v) BSA, and 10% (v/v) fetal bovine serum. Cells were then incubated overnight at 4°C with primary antibodies diluted in PBS containing 1% (w/v) BSA. Following PBS washes, cultures were incubated with Alexa Fluor-conjugated secondary antibodies (Invitrogen; 1:1000) for 1 h at room temperature in PBS containing 1% (w/v) BSA. The cells were then washed with PBS and imaged via confocal microscopy.

The primary antibodies used in these experiments included antibodies specific for tyrosine hydroxylase (Millipore Sigma #AB1542, Burlington, MA; 1:1000), NECAB1 (Thermo Fisher Scientific #PA5-54849; 1:500), ERK1/2 (Thermo Fisher Scientific #13-6200; 1:500), phospho-ERK1/2 (Thr202, Tyr204) (Thermo Fisher Scientific #14-9109-82; 1:500), pS129-aSyn (EP1536Y) (Abcam #ab51253, Cambridge, UK; 1:500], pS129-aSyn (81a) (Millipore Sigma #MABN826; 1:500]), GFAP (Aves labs #GFAP; 1:1000), Iba1 (FUJIFILM Wako Pure Chemical Corporation #019-19741; 1:500), CK1δ (Thermo Fisher Scientific #PA5-32129; #PA5-18509; 1:500), CHMP2B (Thermo Fisher Scientific #12527-1-AP; 1:500), pTau(Ser202/Thr205) (Thermo Fisher Scientific #MN1020; 1:500).

### Sample preparation for proteomic/phosphoproteomic and lipid profiling analyses

Brain tissue samples (∼15-20 mg for rat, ∼10-15 mg for mouse) were resuspended in 300 μL of 25 mM ammonium bicarbonate (ABC) supplemented with protease and phosphatase inhibitors and homogenized in a Precellys Evolution tissue homogenizer using soft-tissue homogenizer CK14 tubes (Bertin Technologies SAS, Montigny-le-Bretonneux, France). The total protein concentration was measured using the bicinchoninic acid (BCA) assay (Thermo Fisher Scientific). For rat brain samples, the Bligh and Dyer (B & D) extraction method (Bligh and Dyer 1959) was performed using lysate volumes equivalent to 25 μg of total protein. The chloroform layer was then carefully transferred to a new tube, dried in a vacuum centrifuge, and used for lipid analysis (see below). Four volumes of cold (−20°C) methanol were then added to the water-methanol layer, and samples were spun at 17,200 x g for 10 min. The supernatant was removed, and pelleted proteins were dried in a vacuum centrifuge and retained for proteomics analysis (described below). For the phosphoproteomics experiments (carried out on both rat and mouse brain samples), tissue lysate containing 500 μg of total protein was precipitated with acetone overnight. The protein pellets (for proteomics and phosphoproteomics experiments) were then resuspended in 10 μL of 8 M urea solution supplemented with 10 mM DTT and incubated at 37°C for 1 h. Next, 10 μL of alkylation reagent mixture (97.5% v/v acetonitrile (ACN), 0.5% v/v triethyl phosphine, 2% v/v iodoethanol) was added, and samples were incubated at 37°C for 1 h. After alkylation, samples were dried in a vacuum centrifuge and resuspended in 80 μL of 0.025 μg/μL trypsin (Thermo Fisher Scientific). The digestion was carried out using a barocycler (50°C, 60 cycles; 50 s at 20 kpsi and 10 s at 14.7 psi). Peptides were then desalted with a C18 silica MicroSpin column (The Nest Group, Ipswich, MA). For phosphoproteomic analysis, phosphopeptides were enriched with PolyMac spin tips (Tymora Analytical, West Lafayette, IN), following the manufacturer’s recommendations.

### LC-MS/MS analysis

Peptide analysis was performed using an UltiMate 3000 RSLC nano HPLC system (Thermo Fisher Scientific) coupled to either a Q-Exactive Orbitrap HF mass spectrometer for rat samples or an Orbitrap Fusion Lumos for mouse samples (Thermo Fisher Scientific). Both LC-MS/MS systems employed Thermo Fisher PepMap trap and analytical columns. Samples were initially loaded onto a PepMap C18 trap column (50 mm × 300 µm, 5 µm particle size, 100 Å pore size) at a flow rate of 5 µL/min for 5 minutes using buffer A (0.1% v/v formic acid in water) to desalt and concentrate peptides. Subsequent chromatographic separation was achieved using an Acclaim PepMap C18 analytical column (500 mm × 75 µm, 2 µm particle size, 100 Å pore size) operated at a flow rate of 150 nL/min. Peptides were eluted over a 160-minute gradient consisting of buffer A and buffer B (0.1% v/v formic acid in 80% v/v acetonitrile in water), as follows: 2% to 27% buffer B over 110 minutes, ramped to 40% over the next 15 minutes, increased to 100% over 10 minutes, held at 100% for 10 minutes, and finally re-equilibrated to 2% buffer B.

For both rat and mouse datasets, full MS (MS1) scans were acquired at a resolution of 60,000 (at m/z 200), followed by MS/MS (MS2) scans at a resolution of 15,000. The scan range was set to m/z 375-1600, with the RF lens set to 30%. The AGC (automatic gain control) target was set to Standard for the Lumos and 3 × 10⁶ for the Q-Exactive HF. Maximum injection times were set to “Auto” (Lumos) or 100 milliseconds (Q-Exactive HF). Data were acquired in data-dependent acquisition (DDA) mode with a cycle time of 3 seconds and a dynamic exclusion duration of 60 seconds to reduce redundant sampling. Peptides were fragmented using higher-energy collisional dissociation (HCD) with 30% normalized collision energy and an isolation window of 1.2 m/z. All instrument parameters and workflows are consistent with previously published protocols.(Glatter, Ludwig et al. 2012, Zembroski, Buhman et al. 2021, Khan, Dutta et al. 2024).

### Lipid profiling analysis

Brain lipids were analyzed using the MRM profiling method (Xie, Ferreira et al. 2021, Reis, Casey et al. 2023). Dried lipid extracts prepared as described above were diluted in injection solvent (acetonitrile/methanol/300 mM ammonium acetate, 3:6.65:0.35 v/v) to obtain a stock solution. This solution was further diluted into injection solvent spiked with 0.1 ng/µL of EquiSPLASH LIPIDOMIX isotopically labeled lipids (Avanti Polar Lipids #330731, Alabaster, AL) for sample injection. MS data were acquired using flow-injection (i.e., without chromatographic separation) of 8 μL of the diluted lipid stock solution delivered using a micro-autosampler (G1377A) to the ESI source of an Agilent 6410 triple quadrupole mass spectrometer (Agilent Technologies, Santa Clara, CA). A capillary pump was connected to the autosampler and operated at a flow rate of 7 μL/min and a pressure of 100 bar. The Agilent 6410 was set to acquire lists of multiple reaction monitoring scans (MRMs) related to lipids. The dwell time for each MRM was 40 ms. The MRMs used were based on the Lipid Maps Standard Database (LMSD) for phosphatidylcholine (PC), phosphatidylethanolamine (PE), phosphatidylinositol (PI), phosphatidylglycerol (PG), phosphatidylserine (PS), ceramides, cholesteryl esters (CE), free fatty acids (just single ion monitoring - SIM), triacylglycerols (TAG), and diacylglycerols (DAG), Cardiolipin (CL). The MRMs related to PC, PE, PG and ceramides were based on protonated parent ions and class-diagnostic product ions. The MRMs related to PS, PI, DAG, TAG, and CE were based on ammonium adducts for the parent ions, class-diagnostic product ions for the PS, PI, and CE lipids, and on fatty acyl neutral losses for the TAG and DAG lipids. Free fatty acids were detected as deprotonated ions in the negative ion mode. The collision energies and fragmentor and cell acceleration voltage settings were determined based on product ion scan experiments using EquiSPLASH LIPIDOMIX isotopically labeled lipids. The electrospray ionization (ESI) source parameters were gas temperature 300°C, gas flow 5.1 L/min, nebulizer 17 psi, and capillary voltage 4 kV for the positive ion mode and 3.5 kV for the negative ion mode.

### Bioinformatics analysis

Raw LC-MS/MS data were processed using the analysis platforms Maxquant (version 2.0.3.1) (Tyanova, Temu et al. 2016) and Perseus (Tyanova, Temu et al. 2016). The data were searched against the *Rattus norvegicus* or *Mus musculus* sequence on UniProt (UniProt 2021) using trypsin as the proteolytic enzyme, allowing up to 2 missed cleavages. Variable modifications were specified for “methionine oxidation” and, in the case of phosphoproteomic analysis, “STY phosphorylation,” while “carbamidomethylation” was assigned as a fixed modification. Other search and data filtering parameters were set as described (Mohallem and Aryal 2024). Raw data files were filtered based on contaminants, and intensity values were log_2_-transformed. Protein/peptide entries present in >70% of the samples were selected for downstream analysis. Data imputation was performed for the replacement of missing values and data quality validation. A two-sample t-test was performed (with a permutation-based false discovery rate (FDR) correction) on samples obtained from PFF-versus monomer-injected animals. Peptide sequences exhibiting a greater than 2-fold increase or decrease in abundance in the brains of animals injected with PFFs versus monomer were subsequently analyzed for gene ontology (GO) enrichment. GO analysis was performed using the Metascape bioinformatics platform (https://metascape.org) (Zhou, Zhou et al. 2019). Phosphoproteins with p < 0.05, q < 0.1, and ≥ 2-fold change were analyzed for protein-protein interactions using STRING. Parameters were set to include all STRING full-network interactions at medium confidence, and MCL clustering was applied to identify natural clusters based on stochastic flow. Heatmaps were generated in MATLAB, and sample clustering based on PCA was performed using the web platform MetaboAnalyst 5.0 (Pang, Chong et al. 2021).

Lipid profiling was performed as described (Xie, Ferreira et al. 2021, Reis, Casey et al. 2023). The MS data were processed using an in-house script to obtain a list of MRM transitions with their respective sums of absolute ion intensities over the acquisition time. Group-average fold changes were analyzed for 13 lipid classes. To determine the group average, intensities were first filtered using a cutoff of 1.3x blank values. Fold change was plotted using Prism software. Statistical analysis was performed utilizing MetaboAnalyst 5.0. Data on relative amounts were autoscaled to obtain a normal distribution, followed by an analysis of heatmap distributions and t-tests with FDR corrections.

### Gene expression quaternary plots

scRNA-seq data were obtained from the Allen Institute for Brain Science Mouse Whole Cortex and Hippocampus 10x dataset using the trimmed means Gene Expression dataset. Gene expression levels (log2 counts per million) were averaged across neurons, astrocytes, microglia, and oligodendrocytes. Proteins identified as significantly differentially phosphorylated in striatally- and cortically-injected mice were plotted according to their expression across these cell types using the R package *plotly*.

### Measurement of proteasomal activity

Proteasomal activity was measured in tissue lysates as described (Berkers, van Leeuwen et al. 2007). Briefly, flash-frozen tissues were lysed in 150 to 400 μL cold assay buffer (50 mM HEPES, pH 7.4, 10 mM NaCl, 1.5 mM MgCl_2_, 1 mM EDTA, 250 mM sucrose, 1 mM freshly prepared ATP) using a 1/8-inch probe tip sonicator (Fisher Scientific, FB120 with CL-18 probe) set to 30% power, with a cycle of 1 s on and 1 s off for 30 cycles. Lysates were then spun at 13,000 × g and 4°C for 20 min, and protein concentrations in the supernatant were determined using a BCA protein assay kit. For activity measurements, proteins (15 to 30 μg) were diluted into 100 or 200 μL of assay buffer containing DTT (1 mM) and 7-amino-4-methyl coumarin (AMC) (25 μM) (Enzo Life Sciences, Farmingdale, NY). Proteasomal activity was quantified by monitoring the rate of fluorescence increase at 37°C using a Synergy fluorescence plate reader (BioTek Instruments, Winooski, VT) with excitation and emission wavelengths set to 360 nm and 480 nm. Control reactions were prepared by supplementing lysates with epoxomicin (Millipore Sigma #324800) at a final concentration of 5 μM to inhibit proteasomal activity. Statistical significance of group differences for each of the 3 enzymatic activities was assessed using one-way ANOVA (Prism software).

### Preparation of brain homogenates and cell lysates for Western blot analysis

Frozen brain samples were homogenized in RIPA buffer (50 mM Tris-HCl, pH 7.4, 150 mM NaCl, 0.1% (w/v) SDS, 0.5% (w/v) sodium deoxycholate, 1% (v/v) Triton X-100) supplemented with protease inhibitors, and the supernatant obtained after centrifugation at 10,000 × g was mixed with Laemmli buffer (Bio-Rad Laboratories, Hercules, CA) containing 5% (v/v) β-mercaptoethanol. Samples were boiled at 95°C for 5 min, separated by SDS-PAGE on 4-20% (w/v) polyacrylamide gradient gels (Bio-Rad), and transferred to 0.4 μm PVDF membranes (Sanyal, Dutta et al. 2019).

For primary neuronal cultures, cells were lysed directly on the culture plate in RIPA buffer supplemented with 100 µM PMSF, phosphatase inhibitor (PhosSTOP, Roche), and protease inhibitor (cOmplete Ultra, Roche). Lysates were collected by scraping and clarified by centrifugation at 16,000 × g for 15 min at 4°C. Protein concentrations were determined using a bicinchoninic acid (BCA) assay, and samples were then mixed with 5x Laemmli buffer and boiled at 95°C for 5 minutes. Equal amounts of protein were separated on 10% (w/v) polyacrylamide gels and transferred to PVDF membranes at 90 V for 2 h in an ice-cold chamber.

For all Western blots, membranes were blocked in either LI-COR blocking buffer (LI-COR Biosciences, Lincoln, NE) or 5% (w/v) BSA in TBS and incubated overnight at 4°C with primary antibody. Primary antibodies were specific for: synapsin (Cell Signaling Technology #D1265, 1:1000), syntaxin (Abcam #ab188583, 1:1000), synaptobrevin (Antibodies Incorporated #1933-SYB, 1:1000), tau (Millipore Sigma #MAB3420, 1:500), β-actin (Millipore Sigma #A5441, 1:2000), ERK1/2 (Thermo Fisher Scientific #13-6200, 1:1000; PhosphoSolutions #500-ERK, 1:1000), phospho-ERK1/2 (Thr202, Tyr204) (Thermo Fisher Scientific #14-9109-82, 1:1000; eBioscience clone MILAN8R, #14-9109-82, 1:750), and β-tubulin (Cell Signaling Technology #15115, 1:1000).

Following washes in TBS containing 0.05% (v/v) Tween-20, membranes were incubated with infrared dye-conjugated secondary antibodies (LI-COR Biosciences) for 90 min at room temperature. Secondary antibodies included IRDye 680RD goat anti-mouse IgG and IRDye 800CW goat anti-rabbit IgG. Membranes were imaged using a LI-COR Odyssey imaging system (Odyssey M Model 3350, LI-COR Biosciences), and band intensities were quantified using ImageJ software (NIH, Bethesda, MD) or LI-COR Image Studio analysis software. Statistical significance was determined using an unpaired t-test in Prism software.

### Golgi-Cox staining

Golgi-Cox staining was performed using the FD Rapid GolgiStain kit (FD Neurotechnologies, Columbia, MD). Brains were immersed in the impregnation solution and incubated for 24 h, after which they were transferred to fresh solution and stored in the dark at 22°C for 2 weeks. The brains were then placed in Solution C and incubated at 4°C for 2 d. To achieve uniform freezing, the brains were immersed in 2-methylbutane cooled to -78°C using a dry ice ethanol bath. Coronal sections (80 μm thick) were then prepared using a cryostat and mounted on gelatin-coated microscope slides. The sections were dried for 2 days, rehydrated in Milli-Q water, and treated with the developing solution. They were then dehydrated in an ethanol series (70%, 80%, and 100% v/v), cleared in xylene, and mounted using DPX mounting medium (Electron Microscopy Sciences). Coverslips were applied to preserve the stained sections for analysis.

### Image acquisition and dendritic spine analysis

Five dendrite sections from layer II/III pyramidal neurons in the cortical region of each mouse were identified and imaged using a laser scanning microscope (Zeiss, Oberkochen, Germany) equipped with a 60x objective lens. High-resolution images were captured from the basal compartment of each neuron and uploaded to Neurolucida 360 software (MBF Bioscience, Williston, VT). For spine density quantification, a dendritic segment of at least 10 μm in length was randomly selected from the basal compartment of each of the five neurons per mouse. Each segment was traced, and dendritic spines were manually counted using Neurolucida 360. Statistical significance of group differences in spine density was assessed using one-way ANOVA (Prism software).

### Cranial window and electrode implant

Under sterile conditions, a custom titanium headplate was attached to the skull using vetbond and metabond dental cement. After headplate implantation, mice were allowed to recover in their home cages for 3 d. Mice underwent habituation of head fixation and running on a 6-in rotating wheel for at least one week before electrophysiological experiments were performed. On the day of the experiment, carprofen and dexamethasone were administered subcutaneously. Following previously established methods (Khan, Dutta et al. 2024), a small craniotomy (< 1 mm) was made over the primary motor cortex (0.4 mm AP, 1.5 mm ML), and the dura was partially removed. A 64-channel electrode (64D Sharp, Mismanidis Lab) was inserted perpendicular to the surface of the pia using a micromanipulator (Sensapex uMp-4, Oulu, Finland). The microelectrode was positioned 1 mm into the motor cortex and allowed to settle for 20 min before recordings began. Signals were digitized at a bandwidth of 0.1 to 10 kHz and sampled at 20 kHz (Intan RHD recording system, Los Angeles, CA). Probe tracks were confirmed histologically after each experiment to verify electrode placement.

### Spike sorting

After the recording session, the electrical signals were processed to sort action potentials from individual neurons. This sorting was accomplished using Kilosort3, followed by a meticulous manual review in Phy2. Neurons were evaluated based on waveform characteristics, stability of firing rates observed throughout the recording session, and autocorrelograms. To analyze the time-varying response of neurons, peri-stimulus time histograms were generated by creating 2000 ms windows during quiescent mouse activity. Average neural activity traces were smoothed by calculating the mean spiking activity across multiple trials and convolving by a Gaussian filter with a full-width at half-maximum (FWHM) of 10 ms. The depth at which each spike occurred was determined based on the electrode that recorded the maximum waveform amplitude. Spiking analysis was conducted on neurons exhibiting sustained activity throughout the entire recording session. Statistical significance of differences in the spiking rates of neurons in the brains of PFF-versus monomer-injected mice was determined using a ranked-sum Wilcoxon test (MATLAB).

### Analysis of spike-spike coherence

To assess the coherence between two spiking signals, we employed discrete signal taper methods as described (Mitra and Pesaran 1999, Jarvis and Mitra 2001, Pesaran, Pezaris et al. 2002, Mitchell, De Houwer et al. 2009). The 2000 ms trial period was segmented into multiple 400 ms intervals with a 200 ms overlap, ensuring full coverage. We removed the DC component from each spike train and applied a single Slepian taper, resulting in an effective smoothing of 2.5 Hz for the 400 ms data windows (NW = 1, K = 1). Confidence intervals were determined using jackknife resampling, involving the exclusion of individual trials. To test whether time-locked trends in firing rate across trials influenced coherence calculations, we performed random shuffles of trial identities and recalculated coherence. This provided a baseline for coherence solely attributed to trends in firing time-locked to the trial. Further, to account for differences in firing rates across neurons within a session, a random number of neurons (5% of the total recorded units) were excluded, and the coherence was recalculated. This was done a total of 3 times to check for biasing introduced by rate-dependent spiking events. Coherence values were computed based on cross-correlations within trials, followed by cross-correlation pooling across trials and normalization by the power spectra of spike trains, which were also pooled over trials.

### Motif enrichment analysis and kinase prediction

Logos for significantly increased or decreased phosphopeptides in binary comparisons were generated using WebLogo (version 3.7.4) (Crooks, Hon et al. 2004). Motif enrichment analysis was done using motifx (R package rmotifx version 1.0) (Chou and Schwartz 2011). Foreground sequences were sequences of total length 15, ± 7 amino acid residues flanking each phosphosite, that were either significantly increased or decreased in binary comparisons (p<0.05, q<0.1). Background sequences were extracted from the rat and mouse proteomes (Uniprot UP000002494 and UP000000589, respectively) using helper packages parseDB and extract Background (R package PTMphinder, version 0.1.0) (Wozniak and Gonzalez 2019). The central residue was set to S, T, or Y, the minimum sequence cut-off was set to 5, and the p-value cut-off was set to 10^-5^. Enriched motifs are represented by their motif score, which is calculated by taking the sum of the negative log probabilities used to fix each position of the motif. Higher motif scores correspond to more specific and statistically significant motifs. Lastly, kinase prediction was performed using NetPhos3.1 online software, with positive predictions having scores >0.5 (Blom, Sicheritz-Ponten et al. 2004).

Kinase activity inference was conducted using RoKAI (rokai.io) (Yilmaz, Ayati et al. 2021). Significantly differentially phosphorylated sites (q<0.1) from all 3 datasets were pooled to generate total unique phosphorylation events. Mouse/Rat UniProt IDs were converted to human IDs using BioMart in R before upload into RoKAI. The uploaded CSV contained 3 columns: “Protein” containing human UniProt IDs, “Position” containing the corresponding human phosphorylation site, and “Quantification” containing phosphorylation site fold-changes. The activity threshold was defined by a minimum number of substrates = 3, with a minimum |Z|-score of 0.5.

## Acknowledgments

This work was supported by a Parkinson’s Foundation Visiting Scholar Award (to S.D.), Purdue Research Foundation Grant (to S.D.), and Bilsland Dissertation Fellowship (to S.D.); funds from the National Institute of General Medical Sciences (T32 GM007752) through the Graduate Training Program in Cellular and Molecular Pharmacology at UCSD (to L.M.R.); National Institutes of Health grants R01AG074273 and R01AG078185 (to X. C.); and grants from the National Institutes of Health (R21NS135424, R01AG098151), the Clinical and Translational Sciences Institute, and Branfman Family Foundation (to J.C.R.).

We thank Dr. Aswathy Chandran, Dr. Andres Salazar-Chaparro, and Dr. Darci J. Trader for assistance with the proteasome assay, Dr. Chandnee Chandrasekaran for assistance with tissue processing, and Dr. Jiaxiang (Jax) Wu and Dr. Maria Olivero-Acosta for guidance and troubleshooting with the Golgi staining protocol and analysis.

## Conflict of Interest

The authors declare no competing financial interests.

## Author contributions

S.D. and J.C.R. conceived the study and prepared the manuscript with input from all authors; S.D. designed and coordinated all experiments, conducted most of the experiments including PFF preparation, *in vivo* surgeries, proteomics data analysis and visualization, IHC, and imaging; J.A.H. assisted with in vivo experiments with rats; A.N.S. and T.J. assisted with *in vivo* experiments with mice and dendritic spine analysis; A.N.S. and L.A.D. assisted with proteomics data analysis and RoKAI kinase analysis; R.M. carried out proteomics sample processing and data acquisition, and assisted with proteomics analysis; L.M.R performed motif enrichment and kinase prediction analysis; H.F.K. performed *in vivo* recording experiments; B.D.C. performed Western blot analyses of rat cortical culture lysates; J.E.G. assisted with dendritic spine analysis; C.R.F. carried out lipid profiling sample processing and data acquisition, and assisted with lipid profiling data analysis; J.F.M. assisted with proteomics sample preparation; X.C. supervised motif enrichment and kinase prediction analysis; K.J. supervised in vivo recording experiments; U.K.A. supervised proteomics sample processing and analysis; L.V.D. supervised PFF preparation and PD modeling in rodents, provided PFF for experiments in rat, and provided input on study design; J.C.R. supervised the study.

## Data Availability

Raw proteomics and lipidomics data are available. The mass spectrometry proteomics data pertaining to this study have been deposited in the ProteomeXchange Consortium via the PRIDE partner repository, with the dataset identifiers PXD057490 (rat phosphoproteome), PXD058091 (mouse phosphoproteome), and PXD057499 (rat global proteome). The lipid profiling data have been deposited to the Purdue University Research Repository (PURR; doi:10.4231/BVVA-4393) and are listed in Supplementary Table 1. A list of differentially expressed proteins (q<0.1) from the rat and mouse phosphoproteomic datasets is provided in Supplementary Table 2.

## Notes

### Competing Interest Statement

The authors have declared no competing interest.

### Summary of Updates

New Figure 7 has been added. Supplementary Figures 2, 8, 12, and 15 have been added New authors have been added, and the order has been changed in the author list.

