## supplementary figures for "Phosphoproteome modifications and cortical circuit dysfunction are linked to the early-stage progression of alpha-synuclein aggregation"

### Supplementary Figure 1

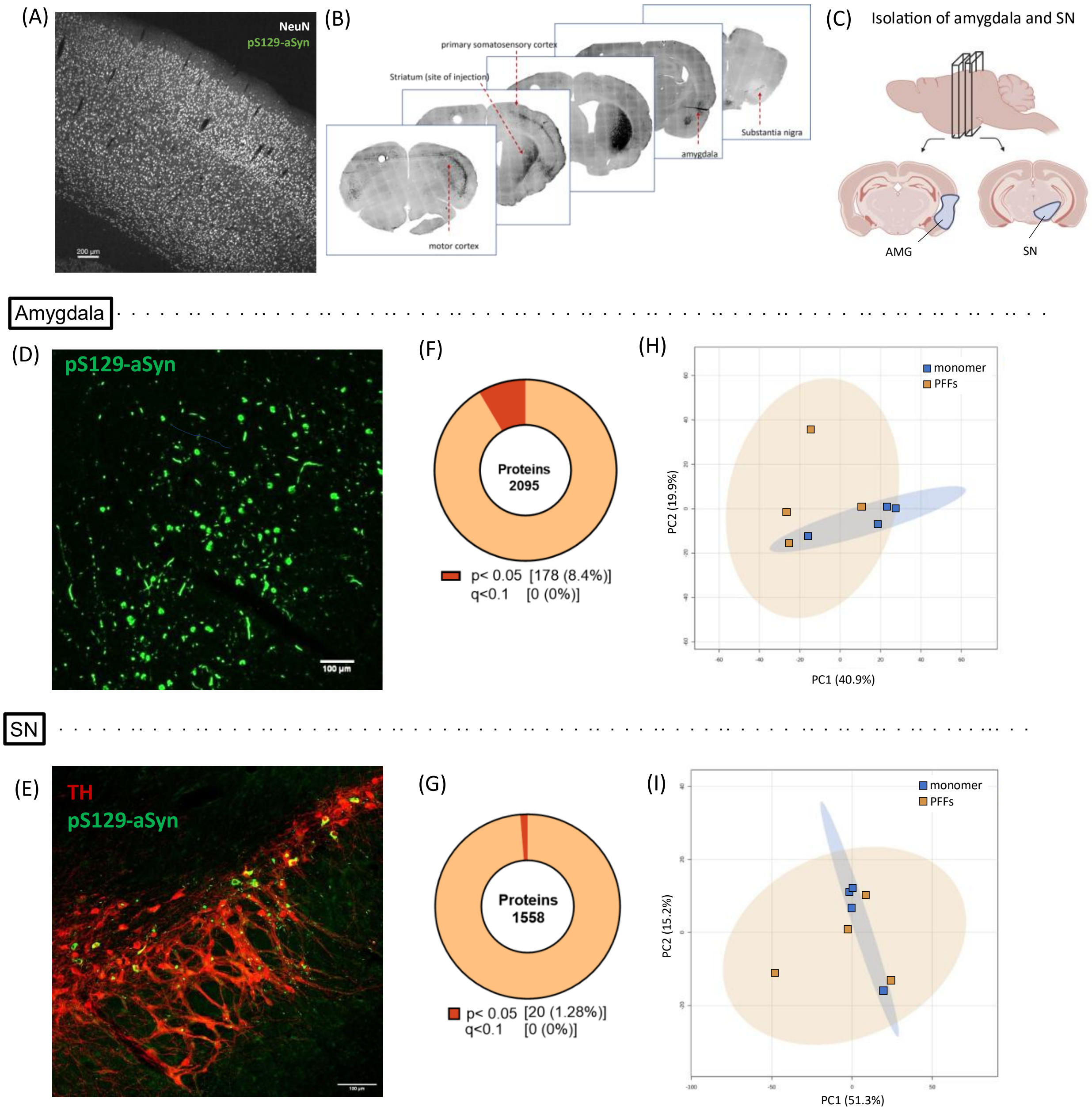

**Supplementary Figure 1: Rats injected intrastriatally with aSyn PFFs do not show evidence of pronounced changes in the global proteome in the amygdala or SN.**

(A) Image showing the absence of pS129-aSyn (81a) signal in a rat cortical section stained for the pan-neuronal marker (NeuN, white) 3 months after monomer injection in the striatum. Scale bar: 200 μm.

(B) Representative images of rat brain sections showing pS129-aSyn immunoreactivity in the cortex, striatum, amygdala, and SN 3 months after aSyn PFF injection in the striatum. Serial coronal brain sections were stained with the EP1536Y antibody.

(C) Schematic of serial brain sections used to collect amygdala (AMG) and SN samples for proteomic analysis (created using https://BioRender.com).

(D-E) Representative images of rat brain sections showing pS129-aSyn immunoreactivity (EP1536Y, green) in the amygdala (D) or SN (E) 3 months after aSyn PFF injection in the striatum. The nigral section in (E) was co-stained for tyrosine hydroxylase (TH, red) to demonstrate the presence of pS129-aSyn inclusions in nigral dopaminergic neurons (n = 3 animals). Scale bar: 100 μm.

(F-G) Pie chart representations of protein hits obtained via global proteomic analysis of homogenates prepared from rat amygdala (F) or SN (G) 3 months after intrastriatal injection with aSyn PFFs or monomer. The chart shows the percentage of hits with $p<0.05$ or $q<0.1$. Each hit was detected in ≥70% of samples in at least one experimental group.

(H-I) Graphs showing the results of unassigned/unsupervised PCA of the $\log_2$-transformed intensities of all protein hits identified in homogenates prepared from rat amygdala (H) or SN (I) as described in (F) and (G).

**Supplementary Figure 2**

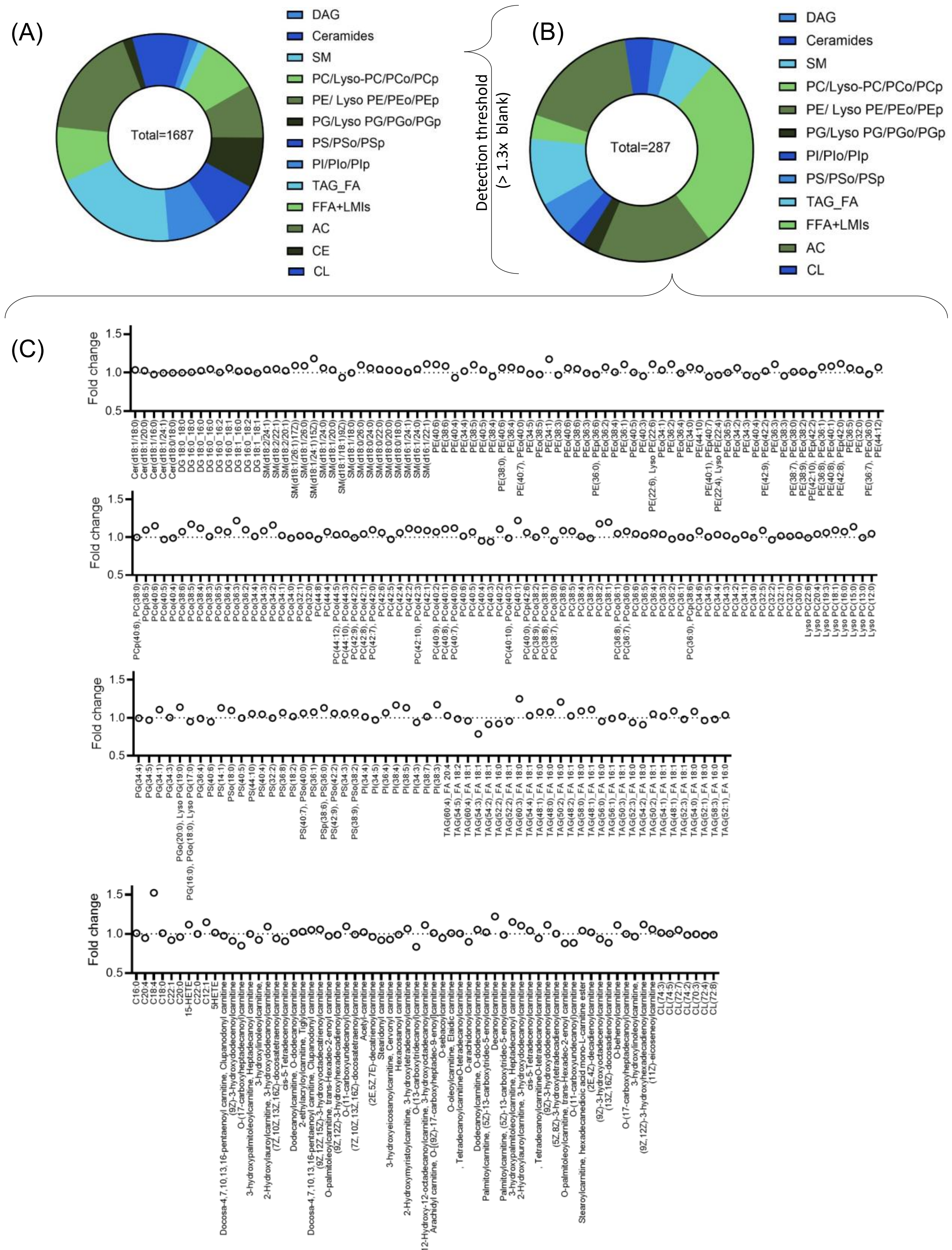

**Supplementary Figure 2: Rats injected intrastriatally with aSyn PFFs do not show evidence of pronounced changes in overall lipid profile. (A)** Pie chart showing the class distribution of all assayed lipid species across 13 lipid classes, including ceramides, diacylglycerols (DAG), phosphatidylcholines (PC and related species: Lyso-PC, PCo, PCp), phosphatidylethanolamines (PE and related species: Lyso-PE, PEo, PEp), phosphatidylglycerols (PG and related species: Lyso-PG, PGo, PGp), phosphatidylinositols (PI, PIo, PIp), phosphatidylserines (PS, PSo, PSp), sphingomyelins (SM), triacylglycerol-derived fatty acids (TAG_FA), free fatty acids and lipid metabolism intermediates (FFA+LMIs), acylcarnitines (AC), cholesteryl esters (CE), and cardiolipin (CL), with total lipid counts indicated. **(B)** Pie chart showing the class distribution of the subset of lipid species (n = 287) that passed a fold-change threshold of ≥1.3 relative to control. **(C)** Dot plot showing fold changes (PFF-injected vs. monomer-injected controls) for the 287 lipid species that met the ≥1.3-fold threshold in cortical homogenates, grouped by lipid class. Each dot represents one individual lipid species.

**Supplementary Figure 3**

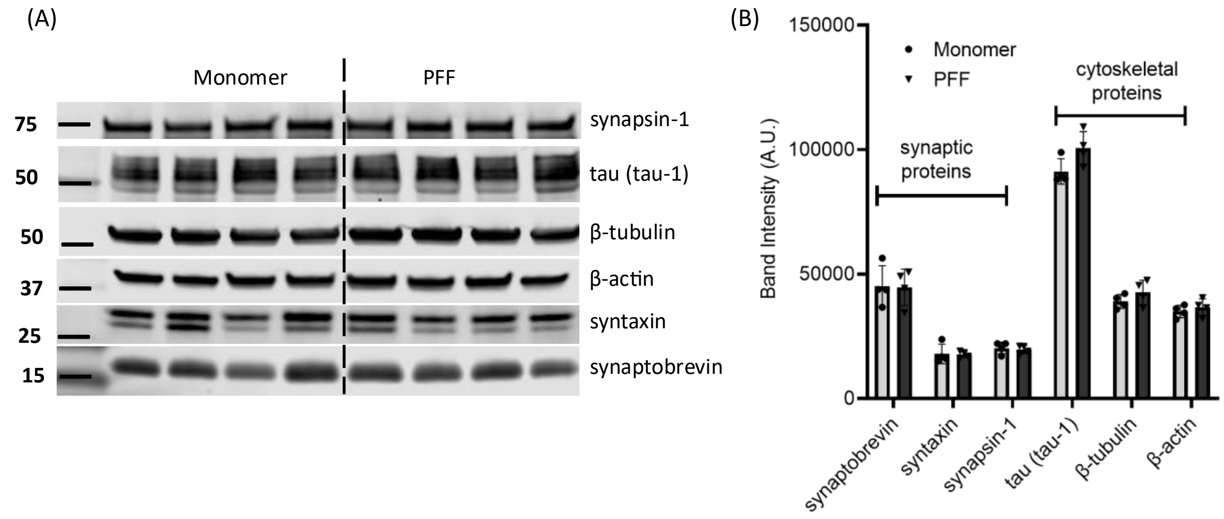


**Supplementary Figure 3: Exposure to aSyn PFFs does not lead to changes in levels of synaptic or cytoskeletal proteins in rat sensorimotor cortex.** (A) Image of Western blot loaded with homogenates prepared from rat sensorimotor cortex 3 months after intrastriatal injection with aSyn PFFs or monomer. The blot was probed with antibodies specific for the synaptic proteins synaptobrevin, syntaxin, and synapsin-1 or the cytoskeletal proteins tau, β-tubulin, and β-actin (n = 4 biological replicates). (B) Graph showing band intensities determined for the Western blot in panel A (mean ± SD). An unpaired t-test revealed no significant differences between the ‘Monomer’ and ‘PFF’ groups for each protein examined on the blot.

**Supplementary Figure 4**

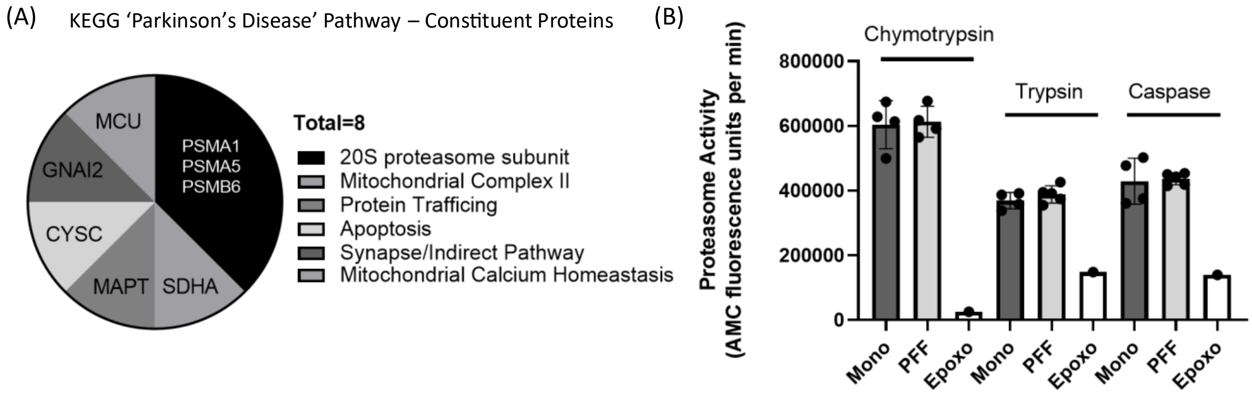


**Supplementary Figure 4: Intrastriatal aSyn PFF injection induces changes in PD-related proteins, including 20S proteasome subunits (without changes in proteasome enzymatic activity), in rat sensorimotor cortex.** (A) KEGG pathway analysis of proteins from the global proteomics dataset with $p < 0.05$ (Figure 1E) identified 'Parkinson's disease' as the most significantly enriched pathway (log $p = -5.17$, log $q = -3.21$). The pie chart displays the 8 hit proteins from our dataset that map to this pathway, grouped by cellular function. The largest subgroup ("20S proteasome subunit") contains 3 of the 8 proteins. PSMA1, proteasome 20S subunit alpha 1; PSMA5, proteasome 20S subunit alpha 5; PSMB6, proteasome 20S subunit beta 6; SDHA, succinate dehydrogenase complex flavoprotein subunit A; MAPT, microtubule-associated protein tau; CYSC, cystatin C; GNAI2, guanine nucleotide-binding protein G(i) subunit alpha-2; MCU, mitochondrial calcium uniporter. (B) Comparison of chymotryptic, tryptic, and caspase-like activities of the 20S proteasome in homogenates from rat sensorimotor cortex 3 months after intrastriatal injection with aSyn monomer or PFFs. Activity was measured using the fluorogenic substrates SUC-LLVY-AMC (chymotrypsin-like), Boc-Leu-Arg-Arg-AMC (trypsin-like), and Ac-Nle-Pro-Nle-Asp-AMC (caspase-like). 'Epoxo' bars represent control homogenates pretreated with the proteasome inhibitor epoxomicin (5 μM, 20 min; n = 1). Data are shown as mean ± SD (n = 4 or 5 biological replicates). One-way ANOVA revealed no significant differences between monomer and PFF groups for any of the three activities.

### Supplementary Figure 5

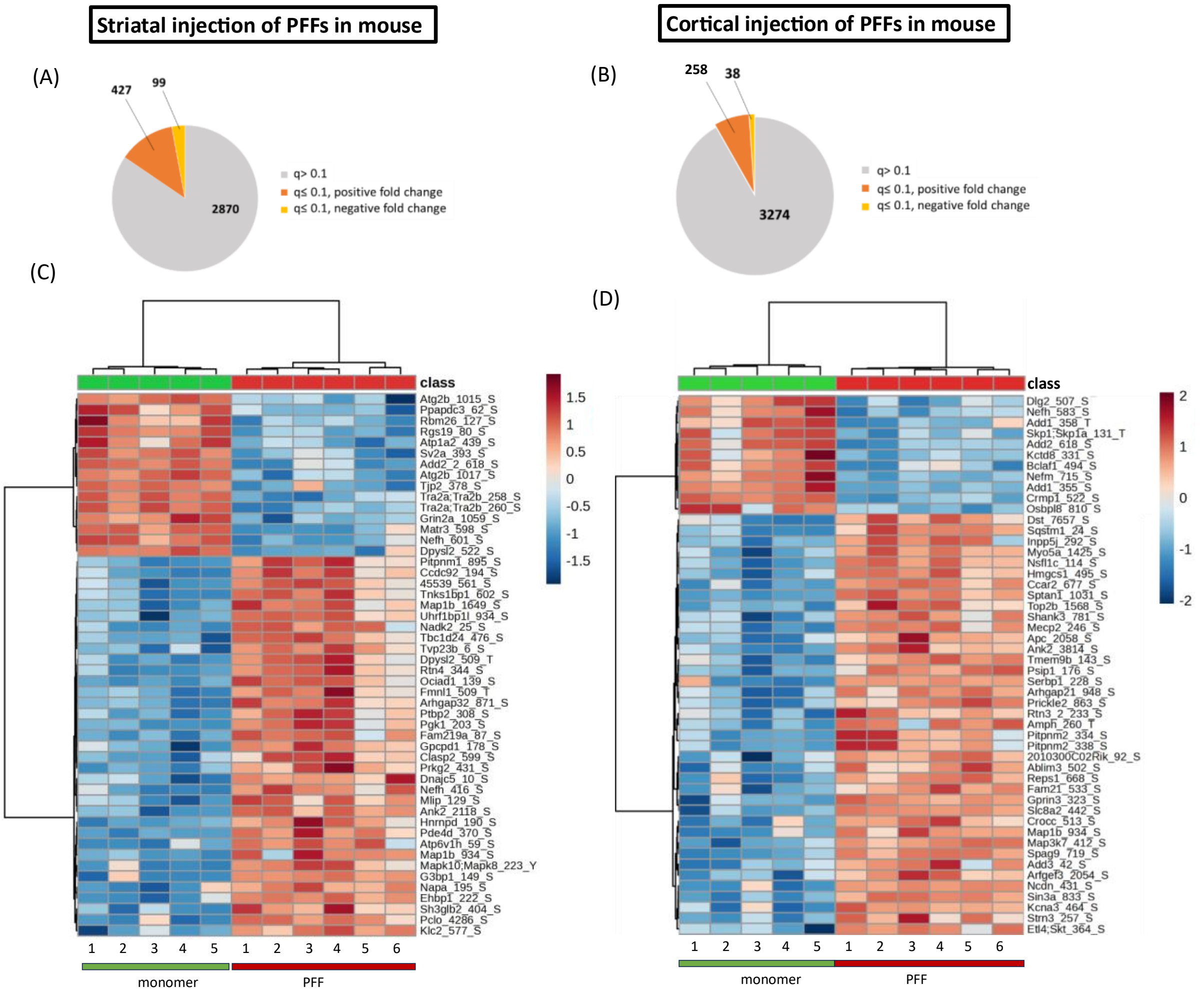

**Supplementary Figure 5: Extended analysis of phosphoproteomic changes in the sensorimotor cortex of mice following intrastriatal or intracortical injections of aSyn PFFs.**

(A, B) Pie chart representations of phosphosite hits obtained via phosphoproteomic analysis of homogenates prepared from mouse sensorimotor cortex 3 months after injection with aSyn PFFs or monomer in the striatum (A) or cortex (B). Each chart shows the percentage of hits with $q<0.1$ with a positive or negative fold change (orange or yellow, respectively). Each hit was detected in ≥70% of samples in at least one experimental group.

(C, D) Clustered heatmaps showing $log_2$-transformed intensities of the top 50 up- or down-regulated phosphosites (i.e., phosphosites with the lowest q-values) in cortical homogenates from mice injected with aSyn PFFs or monomer in the striatum (C) or cortex (D), as described in (A) and (B). Red and blue colors correspond to an increase or decrease (respectively) in phosphosite levels in mice injected with PFFs versus monomer, and the color intensity represents the Z-score-normalized $log_2$(intensity) value. Peptide names are listed as 'protein name_phosphoresidue number.'

**Supplementary Figure 6**

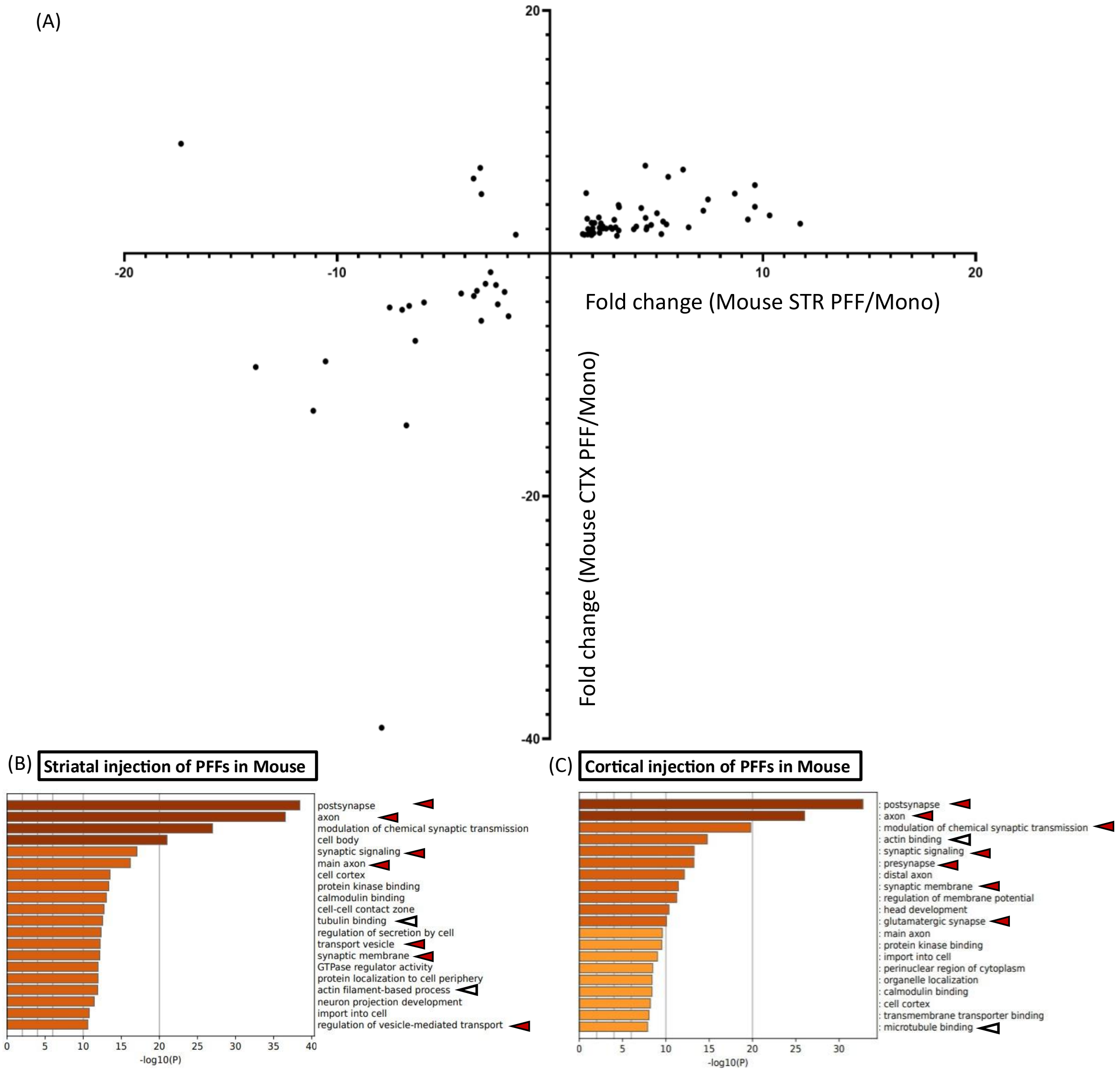


**Supplementary Figure 6: Mice injected with aSyn PFFs in the striatum or cortex show evidence of altered synaptic function and cytoskeletal organization.** (A) Graph depicting fold-change values of significantly altered phosphosites in the sensorimotor cortex of mice injected intrastriatally (STR) or intracortically (CTX) with aSyn PFFs versus monomer. (B, C) Charts showing enriched GO terms in the 'cellular localization' category that represent groups of phosphoproteins in mouse sensorimotor cortex impacted by PFF administration in the striatum (B) or cortex (C), 3 months post-injection. GO analysis was carried out on proteins containing phosphosites with q<0.1 and a fold change of ≥2 (Supplementary Figure 5A,B). Arrowheads highlight items relevant to synaptic transmission (red) or the cytoskeleton (white).

**Supplementary Figure 7**

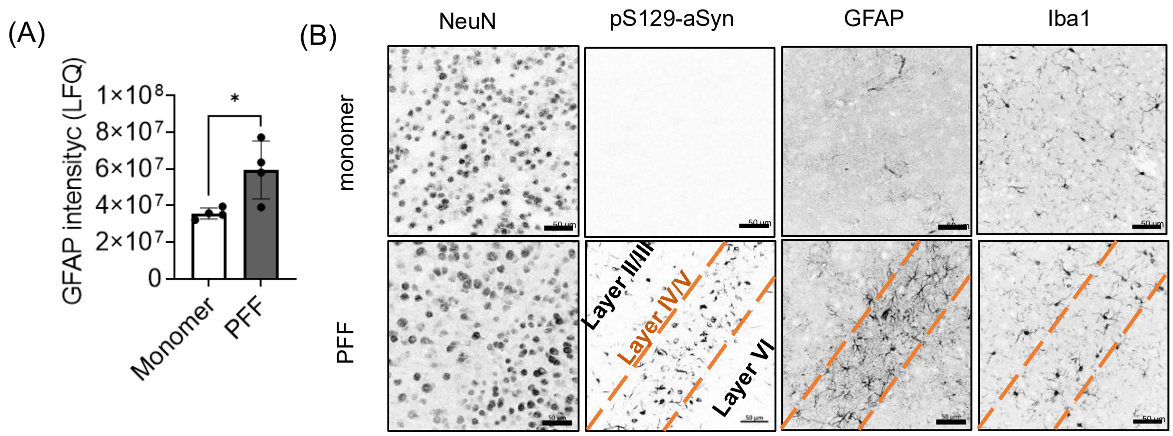


**Supplementary Figure 7: GFAP expression is increased in cortical tissue 3 months post-PFF injection.** (A) Mass spectrometry–derived intensity of the reactive astrocyte marker GFAP in cortical tissue from rats intrastriatally injected with aSyn monomer or PFFs (*$p < 0.05$). (B) Representative IHC images of cortical sections from mice intrastriatally injected with aSyn monomer or PFFs, showing increased GFAP immunoreactivity in PFF-treated animals, particularly in layers IV/V where pathology burden is highest (n = 5 mice). Iba1 immunoreactivity is also elevated in response to PFF injection, albeit to a lesser degree. Staining for the pan-neuronal marker NeuN reveals no significant neuronal loss over the 3-month period. Scale bar: 50 µm.

**Supplementary Figure 8**

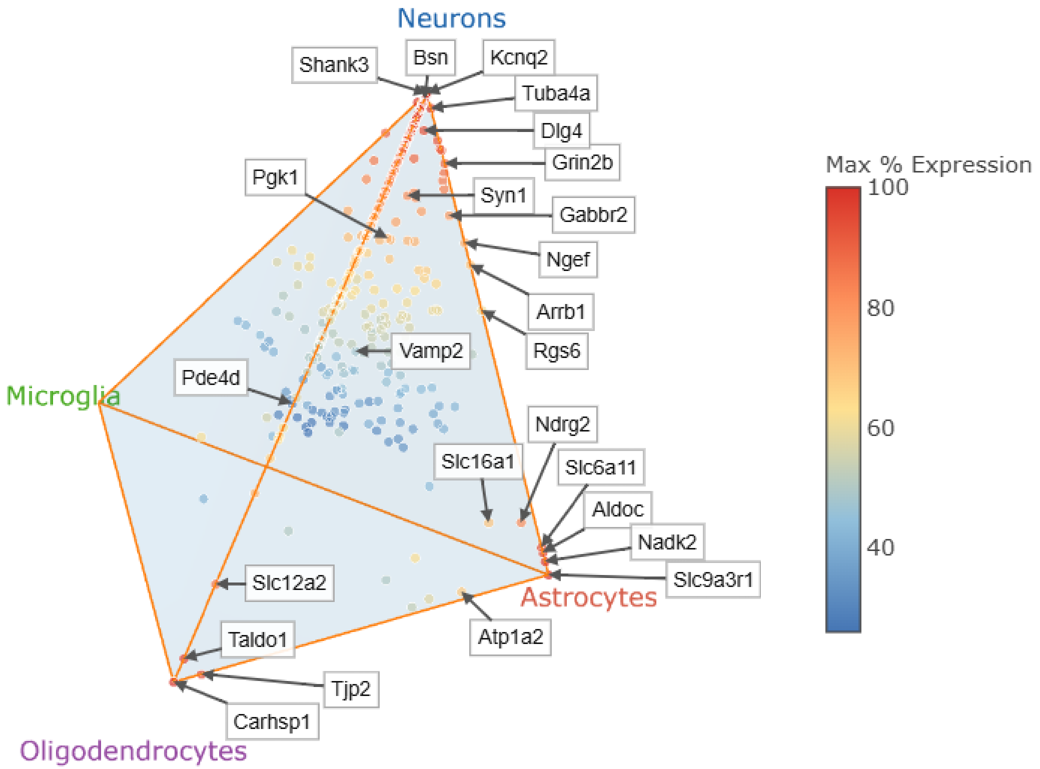


**Supplementary Figure 8: Schematic representation of relative transcriptomic expression levels across neuronal and glial cell types.** Shown are the combined data for differentially phosphorylated proteins identified in the brains of mice injected intrastriatally or intracortically with aSyn PFFs. Each point represents one phosphoprotein and is colored according to its highest percentage of expression in a single cell type (i.e., the cell type in which it is maximally expressed – neurons, astrocytes, microglia, or oligodendrocytes) – see color scale at right. Red points indicate proteins with highly cell-type-specific expression (predominantly expressed in one cell type), whereas blue points represent proteins expressed more broadly across two or more cell types.

**Supplementary Figure 9**

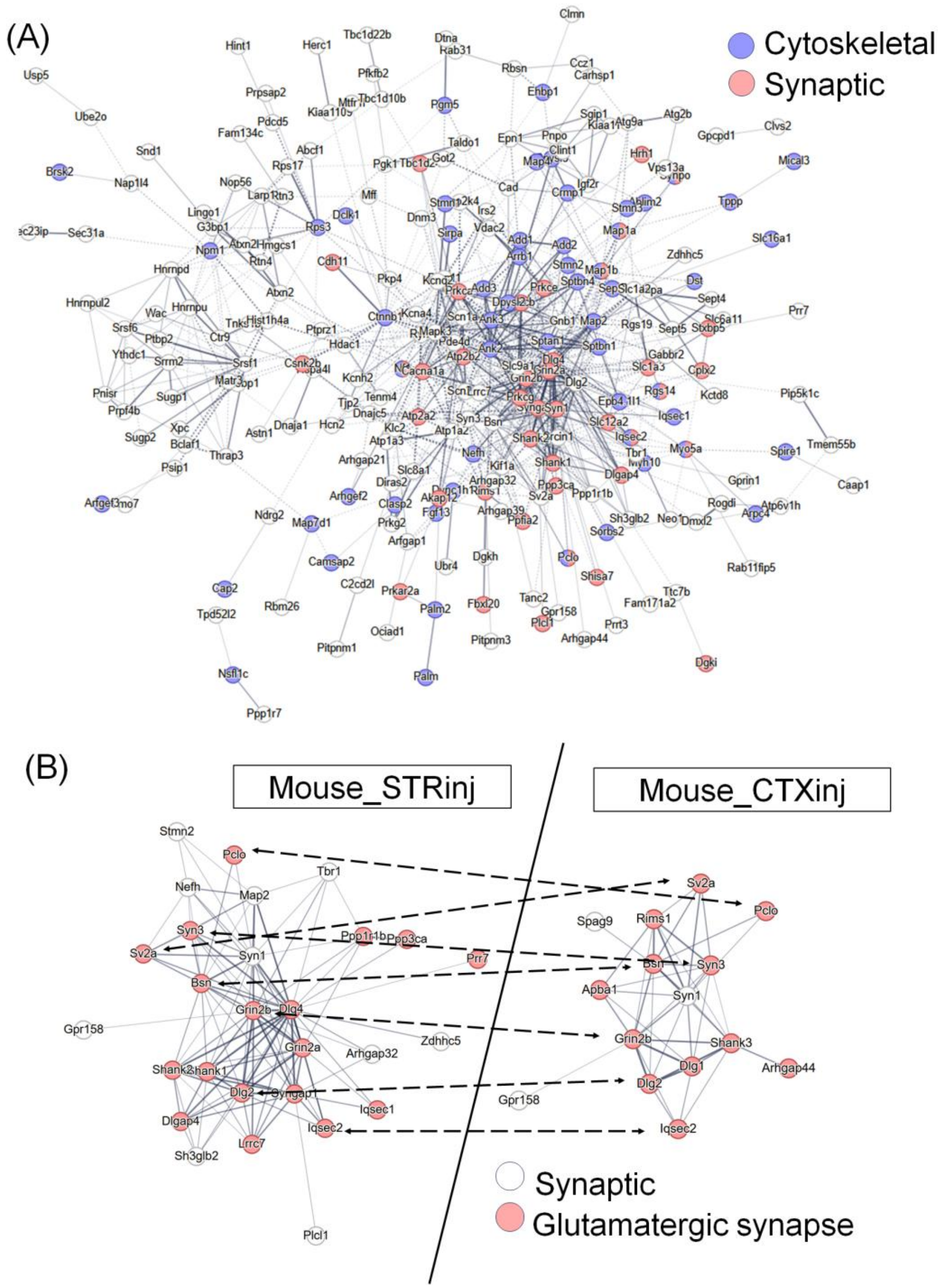

**Supplementary Figure 9: Differentially phosphorylated proteins form interconnected networks enriched for synaptic and cytoskeletal functions.** (A) Network analysis of differentially phosphorylated proteins identified in mice injected intrastriatally with aSyn PFFs revealed extensive protein–protein interactions, forming clusters consistent with the GO terms identified in previous figures, including synaptic and cytoskeletal pathways. (B) Enlargement of the largest cluster, consisting of synapse-related proteins. Proteins involved in excitatory glutamatergic signaling are highlighted in red. The larger cluster size observed in mice injected with PFFs in the striatum versus cortex likely reflects the greater number of input proteins in the striatal dataset (Supplementary Figure 5A,B). Glutamatergic signaling proteins shared between the two datasets are indicated by arrows.

**Supplementary Figure 10**

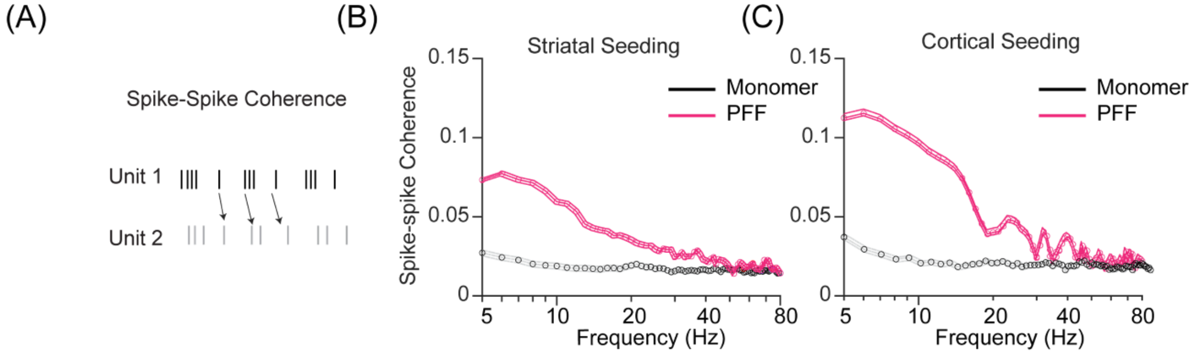


**Supplementary Figure 10: Mice injected with aSyn PFFs in the striatum or cortex show evidence of altered network connectivity.** (A) Schematic of spike-spike coherence where the coherence of two representative units is calculated based on fluctuations of spike rates (see 'Experimental Methods'). (B, C) Graphs showing frequency-resolved paired spike coherence data recorded in mouse sensorimotor cortex 3 months after injection with aSyn PFFs or monomer in the striatum (B) or cortex (C).

**Supplementary Figure 11**

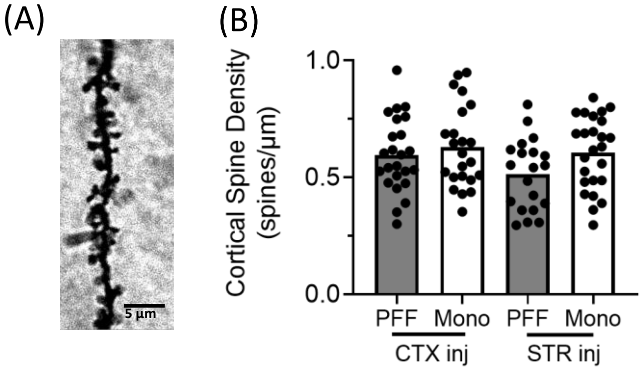


**Supplementary Figure 11: Seeded aSyn aggregation does not lead to a change in cortical spine density in our mouse PFF model.** (A) Representative image of a dendritic section from layer II/III of mouse sensorimotor cortex 3 months after intrastriatal aSyn PFF injection (scale bar: 5 µm). (B) Graph showing the spine density in sections prepared from mouse sensorimotor cortex 3 months after intracortical (CTX) or intrastriatal (STR) injection with aSyn PFFs or monomer. Five dendritic sections were analyzed per mouse, and 4 to 5 mice were analyzed per group. No significant group effects were detected by one-way ANOVA.

**Supplementary Figure 12**

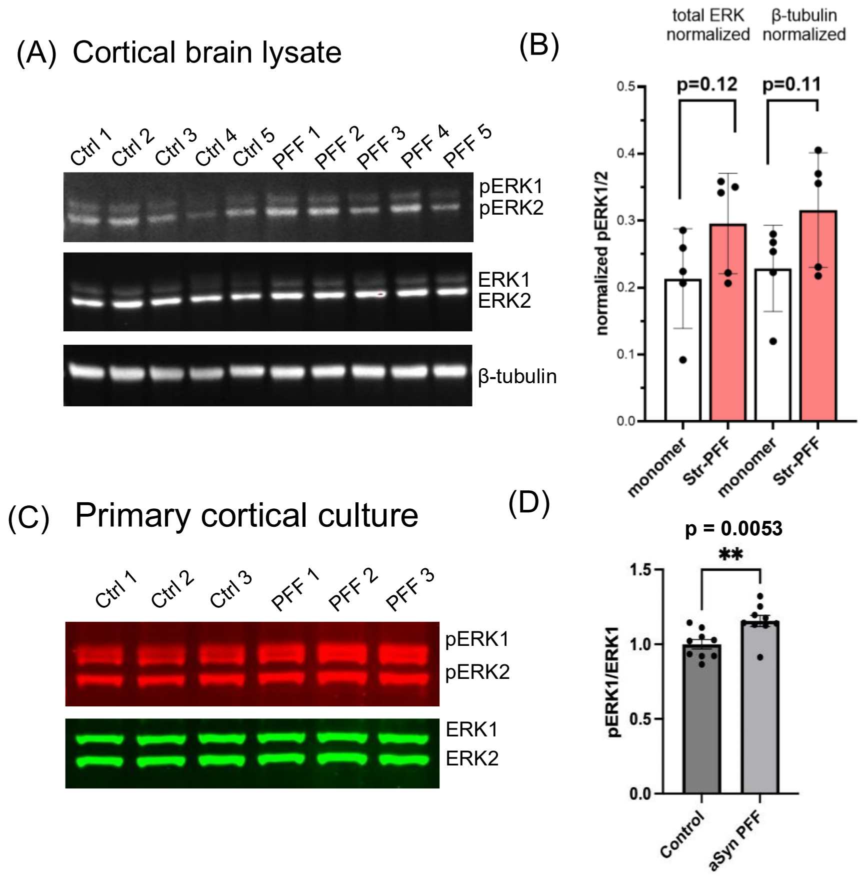


**Supplementary Figure 12: Western blot analyses suggest increased ERK phosphorylation following aSyn PFF exposure.** (A) Representative Western blots of cortical lysates from mice injected with aSyn monomer or PFFs, probed for pERK1/2, total ERK1/2, and β-tubulin. (B) Quantification of pERK1/2 band intensity normalized to either total ERK1/2 or β-tubulin. (C) Representative Western blots of lysates from rat cortical cultures treated with aSyn monomer or PFFs, showing pERK1/2 (red) and total ERK1/2 (green). (D) Quantification of pERK1 band intensity normalized to total ERK1. Data in (B) and (D) are presented as mean ± SEM. Data points in (B) represent independent biological replicates (n=5 mice), with statistical analysis performed using Welch's t-test. Data points in (D) represent technical triplicates from three independent biological replicates (separate cortical dissections). Statistical analysis was performed on log-transformed biological replicate means using Welch’s t-test (n = 3 biological replicates).

**Supplementary Figure 13**

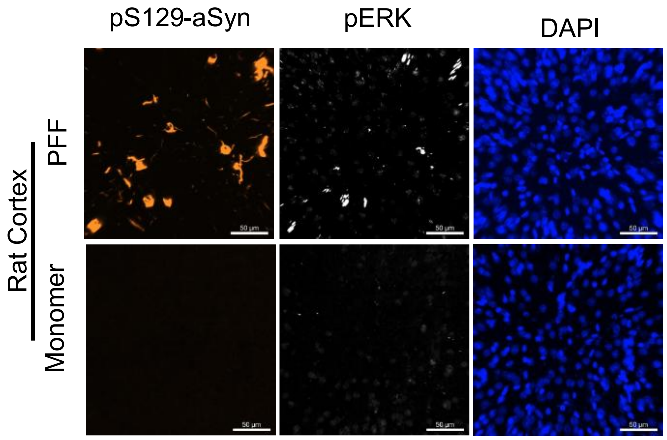


**Supplementary Figure 13: Rats injected with aSyn PFFs exhibit an increase in phospho-ERK levels.** Representative images show increased phospho-ERK (pERK) levels in the sensorimotor cortex of rats injected intrastriatally with aSyn PFFs versus monomer 3 months post-injection (n = 3), confirming ERK pathway activation. Scale bar: 50 μm.

**Supplementary Figure 14**

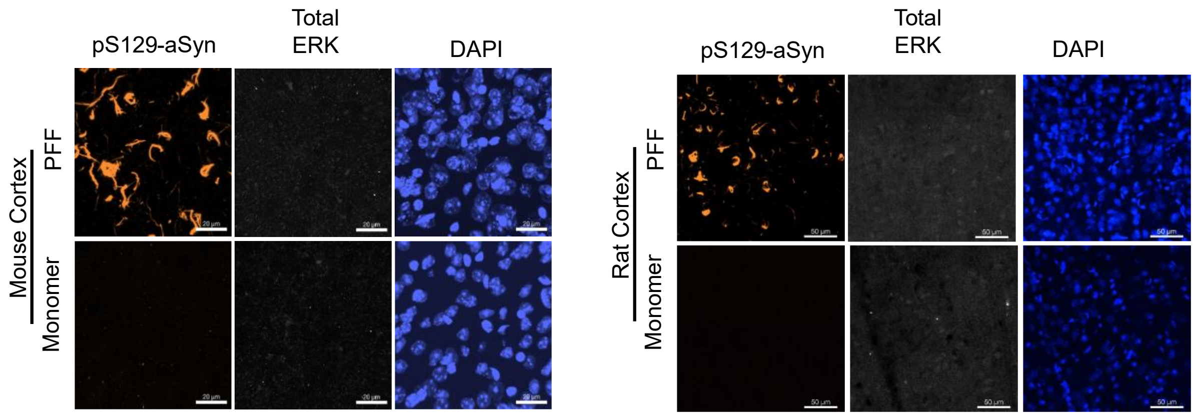


**Supplementary Figure 14: Mice and rats show no evidence of total ERK up-regulation in aSyn aggregate-containing neurons after intrastriatal aSyn PFF injection.** Representative images show no detectable difference in total ERK levels in the sensorimotor cortex of mice (A) or rats (B) injected with aSyn PFFs versus monomer, 3 months post-injection (n = 4). Scale bar: 50 µm.

**Supplementary Figure 15**

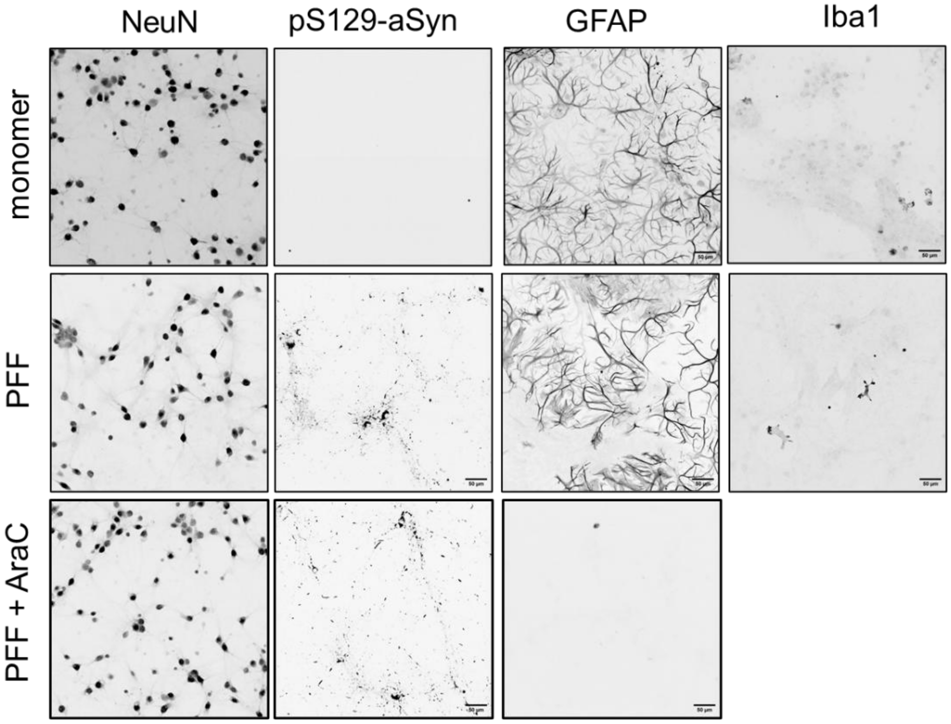


**Supplementary Figure 15. PFF-induced pS129-aSyn aggregation persists following AraC-mediated depletion of astrocytes in primary cortical cultures.** Primary mouse embryonic cortical cultures contained abundant astrocytes with negligible microglial contamination at DIV 10. Treatment with 2 µM AraC at DIV3 markedly reduced the astrocytic population. PFF-induced pS129-aSyn aggregates were readily detected in both untreated and AraC-treated cultures, indicating that aggregate formation was maintained despite substantial astrocyte depletion. Scale bar: 50 µm.
